# An AI-Guided Framework Reveals Conserved Features Governing microRNA Strand Selection

**DOI:** 10.1101/2025.04.30.651563

**Authors:** Dalton Meadows, Hailee Hargis, Amanda Ellis, Heewook Lee, Marco Mangone

## Abstract

MicroRNAs (miRNAs) are central regulators of gene expression, yet how cells choose between the two strands (5p or 3p) of a miRNA duplex during biogenesis remains unresolved. Here, we present a comprehensive, experimentally grounded framework that decodes the logic of miRNA strand selection. Using *Caenorhabditis elegans* as a model system, we developed a high-throughput qPCR platform enabling precise quantification of strand usage across developmental stages and in specific somatic tissues. To uncover the molecular grammar guiding this process, we built a predictive machine learning model trained on experimentally validated strand usage data. This AI-driven model, integrating 77 biologically informed features, accurately predicts strand preference not only in nematodes but also across vertebrates, including humans, revealing compositional and structural biases that are conserved yet functionally repurposed. Our analysis shows that strand selection is not stochastic but follows conserved, context-dependent rules shaped by cellular and developmental cues. To support the research community, we provide open-access resources: a database of strand usage profiles, predictive scores across species, and code and protocols via GitHub. This work offers the first unified, generalizable model for miRNA strand selection, establishing a paradigm that combines large-scale experimentation with AI to reveal a conserved, programmable layer of gene regulation.

## Introduction

miRNAs are ∼22-nucleotide non-coding RNAs that guide the RNA-induced silencing complex (RISC) to target mRNAs, primarily through base pairing within their 3′ untranslated regions (3′UTRs) (1). Once bound, the miRNA-RISC complex (miRISC) typically represses translation of the target mRNA. Both miRNAs and their 3′UTR targets are often conserved across metazoans and play key roles in essential biological processes such as development and morphogenesis.

miRNAs have undergone several pronounced expansions in metazoan genomes (2), coinciding with evolving organismal complexity yet rarely corresponding with increases in protein-coding genes (2). Eukaryotic genomes contain 100-2,000 distinct mature miRNAs in intergenic regions, introns, and in clusters. Following transcription in long primary miRNA transcripts (pri-miRNAs) by RNA polymerase II, they are processed in the nucleus by the microprocessor complex into shorter hairpin structures known as precursor miRNAs (pre-miRNAs) (1). The microprocessor is a multiprotein complex composed of the RNAse III endonuclease Drosha (3), DGCR8/Pasha (4–6) and auxiliary factors. The two strands composing the pre-miRNAs are called the 5p and the 3p strand, respectively. Pre-mRNAs are subsequently exported to the cytoplasm by Exportin-5, where they undergo further cleavage by the enzyme Dicer, producing mature miRNA duplexes (1). Beyond the canonical Drosha-Dicer processing pathway, some miRNAs are generated through non-canonical mechanisms such as mirtron biogenesis (7,8). Mirtrons are short introns that, after splicing and debranching, fold directly into pre-miRNA hairpins and enter the miRNA pathway downstream of Drosha.

The Hsc70-Hsp90 chaperone complex facilitates the ATP-dependent loading of the miRNA duplex into an Argonaute protein, the class of proteins that execute the silencing step (9). This complex is thought to induce conformational changes in Argonaute that enable duplex loading (10–12). Once loaded, one strand of this duplex, the guide strand, is kept within the RISC, while the other, the passenger strand, is typically degraded. Both the 5p and the 3p strands can be selected for miRISC loading and act as guide strands. It is currently unclear how this process, termed ‘miRNA strand selection,’ is executed in detail, which proteins are involved in this decision, and how the guide strand is ultimately chosen between the two strands.

The predominant factors influencing miRNA strand selection are believed to be the specific 5’ nucleotide sequences of each strand and the comparative thermodynamic stability exhibited by the two ends of the miRNA duplex (13). Studies performed using the human Ago2 protein, an Argonaute protein member of the RISC complex, show that the pocket responsible for binding the 5’ nucleotide of the guide strand exhibits a strong preference for Uracil nucleotides (14). This aligns with the prevalent occurrence of uracil as the 5’ nucleotide in miRNA guide strands (2), whereas the passenger strand commonly initiates with a cytosine at its 5’ end.

Furthermore, the miRISC complex exhibits a preference for loading whichever end of the miRNA duplex has lower thermodynamic stability, although not always (15). The initial four nucleotides on each end of the duplex are believed to establish this thermodynamic asymmetry, and in some cases, a slight difference in the number of hydrogen bonds can induce the preferential loading of the less stably paired strand *in vitro* (12).

Consistently, the 5’ ends of guide strands are enriched with purines (2). In comparison, the 5’ ends of passenger strands are predominantly pyrimidine-rich in both human and fly miRNAs, contributing to the thermodynamic asymmetry observed between the two ends of the duplex. In *C. elegans*, single amino acid substitutions within the MID (G553R) and PIWI (G716E, S895F, and S939F) domains of the Argonaute protein ALG-1 invert the strand selection process for specific miRNAs, suggesting a potential role for this protein in the miRNA strand selection process (16). While these mutations play a pivotal role, they do not consistently reverse strand selection for all miRNAs, suggesting that there may be additional internal or external factors involved in this process. Tissue-specific accessory factors and their dosage could also be responsible for miRNA strand selection, altering strand choices based on tissue or developmental contexts, similarly to the role of SR proteins in RNA splicing (17). While thermodynamic and sequence features help explain strand choice, a comprehensive mechanistic framework has been lacking.

*C. elegans* is an ideal model for studying miRNA strand selection due to its compact 3_′_UTRome (18–21), small number of miRNAs (miRbase release 22.1) (22), fully mapped cell lineage (23), and well-characterized transcriptome. Here, we developed a high-throughput qPCR platform to systematically profile miRNA strand selection events across *C. elegans* development and in somatic tissues. These data revealed conserved sequence and structural patterns, which we used to train a machine learning model that predicts strand selection with 83% accuracy and generalizes to human miRNAs.

## Materials and methods

All deionized (DI) water was obtained from either a Purelab Option system (ELGA LabWater) or an Avidity Science system (Duo) and autoclaved before use. Chemicals were ordered from Sigma Aldrich/Merck, VWR, or Thermo Fisher Scientific in molecular biology grade and stored according to company instructions. Gateway^TM^ LR Clonase II Plus^TM^ enzyme mixes, SuperScript^TM^ III Reverse Transcriptase, and PowerTrack^TM^ SYBR Green Master Mix were purchased from Invitrogen. Primers for the HiTmiSS assay were ordered from Life Technologies. Poly(U) Polymerase and restriction enzymes used in validation were ordered from New England Biolabs. The pCFJ150 plasmid was a gift from Erik Jorgensen. The OP50-1 *E. coli*, N2 *C. elegans,* and EG6699 *C. elegans* strains were purchased through the Caenorhabditis Genetics Center.

### Maintenance of *C. elegans* strains

All *C. elegans* strains were cultured on NGM plates seeded with OP50-1 bacteria. All transgenic lines produced for experiments described in this study were maintained at 24°C. When bacteria on each NGM plate had been cleared, *C. elegans* strains were transferred onto new plates via washing into a 15 ml Falcon tube using 15 ml M9 media (22 mM KH_2_PO_4_, 42 mM Na_2_HPO_4_, 85.5 mM NaCl, 1 mM MgSO_4_, sterilized by autoclave) and pelleted for 3 minutes at 400*g*. The M9 media was then aspirated to a volume of 500 ml, and the animals were resuspended via gentle pipetting before being moved onto fresh, seeded NGM plates.

### Synchronization of *C. elegans* strains

The EG6699 strain (24) was maintained at 24°C. To synchronize animals for injection, confluent plates were bleached to isolate the embryos. Each plate was washed into a 15 ml Falcon tube with 15 ml DI water and pelleted for 3 minutes at 400*g*. The water was then aspirated, and the animals were incubated in 5 ml of bleaching solution (0.5 M NaOH, 10% bleach) for 20 minutes with gentle end-over-end rotation. After bleaching, the embryos were washed twice with DI water and moved onto fresh, seeded NGM plates. For use in the HiTmiSS assay, both N2 and *pges-1::GFP::alg-1::unc54 3′ UTR C. elegans* strains were synchronized as described above.

### Primers design for the HiTmiSS Assay

To quantify miRNAs in high throughput, we designed 380 unique forward primers for use in the qPCR amplification step of the HiTmiSS assay. These primers were designed to be complementary to either the 5p strand or the 3p strand of the 190 most expressed miRNAs in the *C. elegans* genome, corresponding to 75% of all *C. elegans* miRNA deposited in miRbase (release 22.1) (25). We did not consider miRNAs whose existence is not supported by broad experimental evidence, nor any miRNAs predicted to exist. A few miRNAs were ignored because their corresponding primers had a very low Tm, making it impossible to process them using our pipeline. All primers were ordered in four 96-well microplates from Life Technologies pre-diluted at a concentration of 50 mM (**Supplemental Table S1**). We designed a universal reverse primer (GCAGGGTCCGAGGTATTC) complementary to the stem-loop sequence of the stem-loop poly(A) primer used to prepare the cDNA for this qPCR reaction (GTCGTATCCAGTGCAGGGTCCGAGGTATTCGCACTGGATACGACAAAAAAAAAAAAA). Twenty-four replicates of this primer were ordered from Life Technologies in a separate array, also at a concentration of 50 mM. Both forward and reverse primers were re-arrayed and diluted to a working concentration of 10 mM immediately prior to use.

### Singletons and Switchers identification

For each strand, we subtracted its relative quantity at one life stage versus the relative quantity of the complementary strand (i.e., for *let-7* 5p, we compared the abundance at the embryo stage versus the abundance of *let-7* 3p at the embryo stage). This was repeated for each life stage at which the expression of that miRNA strand fell above our significance threshold (1 standard deviation above the mean of all quantified miRNAs at a given stage), and any miRNAs for which their expression was greater at each such life stage were deemed *Singletons*. We classified as *5p Singletons* miRNAs where the 5p strand was more expressed at each life stage than its corresponding 3p strand. We classified as *3p Singletons* miRNAs where the 3p strand was more expressed at each life stage than its corresponding 5p strand. We classified as *Switchers* miRNAs that rose above our threshold of significance during at least one life stage, but whose expression levels did not display a consistent pattern of usage of one strand over the other.

### HiTmiSS Assay

To collect sufficient total RNA for analysis of wild type miRNA, confluent NGM plates of N2 strains were synchronized and either washed and pelleted immediately (embryo stage) or incubated at 24°C for 15 hours (larval 1 stage), 25 hours (larval 2 stage), 33 hours (larval 3 stage), 42 hours (larval 4 stage), or 51 hours (young adult stage) before washing and pelleting. All washes were performed in 15 ml Falcon tubes using 15 ml DI water, and pellets were generated by centrifugation for 3 minutes at 400g, resulting in a final pellet volume of 500 μL to 1 mL. Sufficient worms to generate this size pellet differed by life stage, requiring 12 (larval 4 and young adult stages), 18 (larval 2 and larval 3 stages), or 24 (embryo and larval 1 stages) starting confluent plates.

This pellet was then resuspended to a volume of 1.5 ml in ice-cold Triton X-100 lysis buffer (150 mM NaCl, 1.0% Triton X-100, 50 mM Tris-HCl pH 8.0, 0.5 mM EDTA) and transferred to a 2 ml Spectrum tube pre-filled with 0.5 mm zirconium beads. Using a Benchmark BeadBug D1030 microtube homogenizer, each tube was homogenized for 90 seconds, incubated for 2 minutes on ice, then homogenized for an additional 60 seconds. The resultant lysate mixture was centrifuged for 3 minutes using a VWR benchtop mini centrifuge, and the liquid fraction was transferred into a 1.5 ml Eppendorf microcentrifuge tube. To minimize non-aqueous contaminants, this lysate was centrifuged again in the same manner and transferred to a 15 ml Falcon tube pre-chilled on ice. To isolate total RNA, we used a DirectZol RNA MiniPrep Plus kit from Zymo Research, following kit instructions and performing the entire procedure at 4°C. Resultant RNA was eluted in 30 μL nuclease-free (NF) H_2_O.

The protocol for the HiTmiSS assay was adapted from Mei *et al.* (26). We prepared our cDNA from each life stage for qPCR analysis via polyuridinylation using Poly(U) Polymerase (NEB) as per manufacturer instructions followed by a 20-minute heat inactivation at 65°C. We performed a first-strand reaction on this polyuridinylated RNA using Superscript III Reverse Transcriptase and stem-loop poly(A) primers under recommended conditions (Invitrogen). We then performed qPCR in high throughput using this cDNA, SYBR Green master mix (Invitrogen), and our primers as described above. We used Applied Biosystems QuantStudio 3 and QuantStudio 7 qPCR machines to perform the screen. For each life stage, four 96-well plates were prepared with 40 μL reactions in each well, along with a positive control reaction containing 10 pg pDONR 221 ROG instead of cDNA and the necessary forward and reverse primers to amplify *gfp*. Each plate was then re-arrayed in triplicate into a 384-well plate using a Biomek FX^P^ Laboratory Automation Workstation for a total of three 10 μL reactions per miRNA strand per life stage. An additional plate was prepared with no cDNA or plasmid spike-in DNA as a non-template control. These qPCR reactions were carried out under standard SYBR Green reagent conditions on an Applied Biosystems QuantStudio 7. Relative miRNA levels were calculated using the 2^-ΔΔCt^ method (see below), and normalized using the spike-in positive control. These results were then plotted logarithmically as a heatmap, with data below our analytical threshold visualized in black. Data were ordered by type (*5p Singleton*, *3p Singleton*, or *Switcher*) and visualized using the Heatmapper2 web application (27).

### Additional MIQE-Compliance-related methods

#### RNA Integrity Reporting

RNA purity and integrity were assessed using a Nanodrop spectrophotometer (Thermo Scientific). All samples had 260/280 ratios between 1.9-2.1 and 260/230 ratios >1.8.

#### Primer Validation

Primer specificity was inferred from the absence of amplification in non-template controls and the presence of a single, well-defined amplification curve for each reaction. All primer sets produced consistent Ct values across independent replicates and developmental stages, and no secondary amplification products were detected in NTC reactions. The HiTmiSS assay is a high-throughput adaptation of Mei et al. (26), who demonstrated primer specificity using stem-loop qPCR with strand-specific forward primers. Because our assay uses identical primer architecture, specificity is expected from design and was verified by negative controls.

#### Biological Replicates

For each developmental stage and tissue, RNA was extracted from independent biological populations of worms cultured on separate plates. Each qPCR was run in technical triplicate.

#### NTC Controls

Non-template controls did not produce detectable amplification across all primer sets.

#### Reference Stability

*let-7* and *lin-4* were chosen as reference miRNAs based on their stable expression across developmental stages and tissues (confirmed by low Ct variance across samples).

### Cycle Threshold Analysis of the HiTmiSS Assay

HiTmiSS assay queries each miRNA strand separately, normalized by a per-assay positive and negative control. The qPCR results containing the cycle threshold (Ct) values for each tested miRNA were exported in Microsoft Excel 2019 format. A comprehensive list of these data can be found in **Supplemental Table S2**. Ct values were averaged across experimental replicates, then subjected to a 2^-ΔΔCt^ analysis to calculate initial amounts of each miRNA in the sample, consistent with experimental standard (28). The Ct value for any given miRNA strand was subtracted from the Ct value of the non-template control (a qPCR reaction lacking any cDNA) for that miRNA strand. This was termed the ΔCt for that miRNA. The same was done for the positive control for that sample. Then, for each sample, this difference was subtracted from the difference obtained for that positive control, normalizing samples to the same positive control and giving us the ΔΔCt for that miRNA strand. 5p/3p comparisons for the same pre-miRNA, use a universal reverse primer and strand-specific forward primers matched for Tm, length, GC content, and amplicon size. The inverse log base 2 of that ΔΔCt was then calculated to approximate the initial amount of that miRNA strand present in the cDNA relative to the positive control. This was deemed the 2^-ΔΔCt^ value for that miRNA strand. To determine whether a miRNA was considered present in a sample, we decided on a strict threshold of one standard deviation above the mean of all 2^-ΔΔCt^ values for that stage of development. MiRNAs quantified at levels above this threshold were considered significant for the purposes of this study.

### HiTmiSS Assay in the Intestinal Tissue

To isolate tissue for use in an intestine-specific HiTmiSS assay, we first performed protein immunoprecipitation followed by miRNA isolation. The fluorescent *pges1::gfp::alg-1::unc54 3′UTR C. elegans* strain previously prepared by our lab (29) was grown to confluence, synchronized, and pelleted as described above for each life stage, concurrent with N2 *C. elegans* as a negative control.

Each pellet was resuspended to 1.5 μl using our Triton X-100 lysis buffer with added protease inhibitor (two Roche cOmplete ULTRA mini EDTA-free protease inhibitor tablets per 50 mL lysis buffer) and transferred to a zirconium bead tube to be homogenized as previously described. During this homogenization step, anti-GFP magnetic antibody beads (Chromotek GFP-Trap Magnetic Agarose) were prepared by diluting 25 μl bead slurry in 500 μl dilution buffer (150 mM NaCl, 50mM Tris-HCl pH 8.0, 0.5 mM EDTA) and rotating end-over-end at 4°C for 3 minutes.

The beads were pelleted using a benchtop centrifuge, and the buffer was aspirated using a magnetic tube rack. 500 μl dilution buffer was added again, and this dilution process was repeated twice. We then recovered aqueous lysate from the homogenized mixture as previously described and diluted it using 300 μl dilution buffer. This diluted lysate was transferred to a tube containing the prepared anti-GFP beads and left to rotate end-over-end at 4°C overnight. To clear unbound lysate, the beads were pelleted using a benchtop centrifuge, and the buffer was aspirated using a magnetic tube rack. 500 μl of wash buffer (150 mM NaCl, 50 mM Tris-HCl pH 8.0, 0.1% Triton X-100, 0.5 mM EDTA) was added, the beads were pelleted, and the buffer was aspirated using a magnetic tube rack. This wash step was repeated five additional times. The beads were then resuspended in 100 μl wash buffer and RNA was isolated using a DirectZol RNA MiniPrep Plus kit from Zymo Research, following kit instructions and performing the entire procedure at 4°C. Resultant RNA was eluted in 30 μL NF H_2_O.

The HiTmiSS assay and subsequent analysis was performed as described above, with the following modifications: (1) Forward primers were re-arrayed from the 96-well microplates described above, resulting in a screen of the 93 most highly quantified miRNAs (93 5p and 93 3p strands) from our whole worm analysis. This was done to avoid redundant quantification of miRNAs which in whole worm fell below our quantification threshold and were therefore unlikely to display significant expression in a tissue-specific screen performed on immunoprecipitated miRNAs. (2) Only embryo (E) and young adult (YA) stages were included in this analysis. (3) The analytical threshold per life stage was set at the mean expression of all miRNA strands at that life stage, due to globally decreased cycle threshold values detected in a single tissue relative to our spike-in positive control used for normalization. (4) Data was ordered by overall strand expression across both life stages, leading to only two major clusters on the resultant heatmap. Additionally, miRNAs which fell below our analytical threshold for both strands at each life stage were excluded from the heatmap for clarity.

### qPCR of miRNAs targeting overexpressed intestinal 3**′**UTRs

We used two *C. elegans* strains previously generated and published by our lab to measure miRNA strand levels upon intestinal overexpression of their target 3′UTRs,. (29). Each expressed the promoter for intestine-specific gene *ges-1* fused to a dual color ROG construct (a gene encoding *mCherry* and *gfp* separated by a trans-spliceable operon) fused to the 3′ UTR of either *fox-1* or *hrp-2* (*29*). Each of these strains was grown to confluence in 12 NGM plates in mixed stage and subjected to quantification using the same approach as the HiTmiSS assay described above, though rather than quantifying every miRNA in these animals, we quantified the 5p and 3p strands of *miR-54*, *miR-60*, *miR-232*, and *miR-239b*. These quantifications were carried out in 96-well plates using cDNA (both prepared from our strains as well as control cDNA gathered from N2 *C. elegans*), SYBR Green master mix, and the appropriate forward and universal reverse primers for the miRNAs of interest, re-arrayed from our standard HiTmiSS plates. Each 96-well plate contained two positive controls in *let-7* and *lin-4*, as well as a non-template negative control for each set of primers. These qPCRs were carried out in experimental triplicate under standard SYBR Green reagent conditions on an Applied Biosystems QuantStudio 3.

Relative miRNA levels were calculated using the 2^-ΔΔCt^ method as described above and normalized using the average of *let-7* and *lin-4* expression for each sample. These results were then averaged across two replicates and analyzed using Microsoft Excel 2019. Data were visualized as log_2_ of the change in expression from wild-type N2 negative control to either of our experimental strains. The threshold of significance was set at one as per standard analytical procedure.

### Comparison of HiTmiSS vs RNA-seq

To compare HiTmiSS results with RNA-seq data, we retrieved small RNA sequencing datasets from Kato et al. (30) and Stubna et al (31) via the Sequence Reads Archive (SRA). Both studies employed deep sequencing (Illumina) to profile the expression of small RNA and miRNA across *C. elegans* developmental stages. For Kato et al., we analyzed the following datasets: SRR014966 (embryo), SRR014967 (L1), SRR014968 (L2), SRR014969 (L3), SRR014970 (L4), and SRR014971 (young adult). For Stubna et al., we analyzed the following datasets: L1: SRR29013898 (rep1), and SRR29013897 (rep2). L2: SRR29013894 (rep1), and SRR29013893 (rep2). L3: SRR29013890 (rep1), and SRR29013889 (rep2). L4: SRR29013889 (rep1), and SRR29013885 (rep2). YA: SRR29013882 (rep1), and SRR29013881 (rep2). Early Embryo: SRR29013881 (rep1), and SRR29013905 (rep2). Late Embryo: SRR29013902 (rep1) and SRR29013901 (rep2). The early and late embryos were pooled together and averaged for the analysis described below. To assess both the presence of miRNA and its strand usage, we developed a custom analysis pipeline. Sequencing reads were first converted to FASTA format using the FASTX-Toolkit (http://hannonlab.cshl.edu/fastx_toolkit/), then aligned using BLAT (32) against a reference database containing all known *C. elegans* miRNA strands (miRbase) (25) in FASTA format, with the following parameters: /blat -t=dna - q=dna -tileSize=6 -minIdentity=10 -minMatch=1 -minScore=10 stage_specific.fa stage_specific.psl. To increase stringency, we filtered the resulting PSL files to retain only matches with strand = “+”, and with Q_start = 0 (for Kato et al.) and 100% match (For Stubna et al.,) then quantified the occurrence of each miRNA strand at each developmental stage, with at least 50 reads. For the Kato et al. dataset, miRNA strands with ≥50 reads were considered for analysis. The numbers of unique 5p and 3p strands, together with their corresponding median read counts, were as follows: Embryo, 158 (5p) and 147 (3p), totaling 305 unique miRNA strands (median = 139.5 reads); L1, 81 (5p) and 87 (3p), totaling 168 unique miRNA strands (median = 359 reads); L2, 85 (5p) and 82 (3p), totaling 167 unique miRNA strands (median = 373.5 reads); L3, 74 (5p) and 74 (3p), totaling 148 unique miRNA strands (median = 483 reads); L4, 67 (5p) and 81 (3p), totaling 148 unique miRNA strands (median = 589.5 reads); and YA, 70 (5p) and 82 (3p), totaling 152 unique miRNA strands (median = 507 reads). For the Stubna et al. dataset, a consistent set of 192 (5p) and 190 (3p) strands was identified across all stages. The median read counts per replicate were as follows: Early embryo, 6,501 (rep1) and 5,089 (rep2); Late embryo, 7,805 (rep1) and 7,630 (rep2); L1, 5,362 (rep1) and 6,210 (rep2); L2, 5,310 (rep1) and 7,390 (rep2); L3, 9,187 (rep1) and 8,782 (rep2); L4, 11,605 (rep1) and 11,313 (rep2); and YA, 16,194 (rep1) and 30,042 (rep2). We then used these counts to calculate the expression value of each miRNA strand normalized to each developmental stage.

### Structural Analysis of miRNAs

The experimentally validated *5p* and *3p Singletons* were obtained using our HiTmiSS assay. Each entry included the entire hairpin sequence and its predicted secondary structure in dot-bracket notation (33) (**Supplemental Table S2)**. To focus specifically on mature miRNA regions, only the sequences of each mature miRNA duplex were used for all downstream analyses. Since the *5p* and *3p Singleton* datasets contained different numbers of miRNAs, we balanced them before analysis by undersampling the larger group to match the sample size of the smaller class, ensuring unbiased statistical comparison between 5p and 3p strands.

We extracted 77 numerical features from each *5p* and *3p Singleton* (**Supplemental Table S3)**. The extracted features included: Base composition (fraction of A, U, G, C), motif presence (e.g. UGU, GAG, CCU), positional characteristics (first/last nucleotide identity), dinucleotide frequencies (e.g., di_GG, di_CC), GC content and GC-related asymmetry metrics (GC_diff, GC_ratio_5p3p), strand lengths, and number of mismatches (unpaired positions in the dot-bracket structure). The full list of the features and their descriptions is shown in **Supplemental Table S3**.

Feature calculations were implemented in Python using regular expressions and sequence statistics functions. Structural mismatch counts were determined by counting unpaired characters within the uppercase-aligned mature duplex. Comparative statistical analysis between 5p and 3p strands was conducted using Welch’s two-sample t-test. All features were tested independently, and results were ranked by p-value. Features with p < 0.05 were considered statistically significant. Bar plots were generated for each significant feature using the mean ± standard deviation across strands.

### miRNA strand selection ML model

To predict dominant strand selection (5p vs. 3p) in miRNA hairpins, we utilized ChatGPT GPT-4o (OpenAI) to assist in developing and training a Random Forest classifier based on 77 biologically curated features, organized into functional categories capturing distinct structural and sequence-related aspects of miRNA hairpins. These features capture thermodynamic stability, base-pairing patterns, and positional relationships within the miRNA hairpin. The model was trained on our validated *5p* and *3p Singletons* identified using our HiTmiSS assay, and feature values were computed using custom Python scripts that parse dot-bracket RNA structures and extract both absolute and relative measures for each strand. The weights of the features were optimized using known *Singletons*. The final model incorporates the following features, divided into five categories. The first category is the *Dinucleotide Composition* features, which quantify the frequency of specific two-nucleotide motifs within the 5p and 3p regions, capturing local sequence patterns that can influence strand selection and structural flexibility. The second category is the *First/Last Nucleotide Bias* features, which record the identity of the first and last nucleotides in each strand, reflecting known preferences in Argonaute loading where terminal nucleotides strongly impact strand choice. The third category is the *GC Content and Entropy* features, which assess the global nucleotide composition and thermodynamic diversity within each strand, including overall GC percentages, GC ratios between strands, and Shannon entropy measures that reflect structural variability and rigidity. The fourth category is the *Base-pairing Properties* features, which describe the fraction of nucleotides engaged in base pairing within the 5p and 3p regions, providing a measure of local hairpin stability and asymmetry. Finally, the *General Structural* features, which include the overall hairpin length and the length difference between the 5p and 3p strands, capture broader geometrical properties that may indirectly affect miRNA maturation and strand retention. The complete set of features, their description, and the weight used are shown in **Supplemental Table S3**. The model showed high performance in distinguishing 5p and 3p usage from a pool of known *Singletons* as a test dataset (*5p Singleton*: accuracy 100%, precision 75%, recall 100%; *3p Singleton:* 82% accuracy, 100% precision, 82% recall). The ML model uses a Random Forest classifier implemented in scikit-learn (v1.3.1). Model hyperparameters were optimized via grid search with 5-fold stratified cross-validation, tuning: n_estimators: [100, 250, 500], max_depth: [5, 10, None], min_samples_split: [2, 5, 10], class_weight: [‘balanced’, None]. The final selected parameters were: RandomForestClassifier (n_estimators=250, max_depth=None, min_samples_split=2, class_weight=’balanced’). To prevent overfitting and enhance interpretability, recursive feature elimination (RFE) with cross-validation was used to rank features by importance. Features with near-zero variance and multicollinearity (|Pearson’s r| > 0.9) were removed. The model consistently retained ∼45–55 features, depending on the species subset. Performance was assessed on independent held-out test sets not used during training. Metrics included: Accuracy: 0.85 (*C. elegans*), 0.83 (human), Precision (5p/3p): 0.87 / 0.80, Recall (5p/3p): 0.89 / 0.78, F1 Score (5p/3p): 0.88 / 0.79, AUROC: 0.91 (*C. elegans*), 0.89 (human). Class balance was maintained using SMOTE oversampling to correct for the slight enrichment of 5p strands in the source datasets. We confirmed that the model generalizes well by testing it on separate datasets (not used during training or tuning), including across species (e.g., training in *C. elegans*, testing in humans).

### Unsupervised benchmark of strand-selection predictors

We evaluated three predictors on the same set of human miRNA duplexes (n = 854): (i) HiTmiSS (77-feature RF; outputs 5p/3p and P(5p)); (ii) a 5′-nucleotide rule (guide predicted by first-base preference U > A > C > G; ties to 5p); and (iii) an end-stability proxy (ΔΔG) that scores the first 5 nucleotides at each arm’s 5′ end against the reverse-complement of the opposite arm (GC=3, AU=2, GU=1, mismatch=0), predicting the arm with the less-stable 5′ end. Because a comprehensive gold standard is unavailable, we tested the three predictions using the Dawid-Skene label-aggregation model, which jointly infers (a) a latent “true” label for each duplex from the pattern of agreements and (b) a confusion matrix (reliability) for each predictor. We report each method’s estimated accuracy as the prior-weighted proportion of correct calls from its inferred confusion matrix. All parameters were prespecified (no tuning), and the same items (n=854) were used for all methods.

### ML model predictions in human miRNAs

Human miRNA expression data were obtained from a harmonized dataset from the human miRNA tissue atlas (34) containing strand-specific Reads Per Million Mapped (RPMM) values across 65 healthy tissues and cell lines of human origins. Each sample contained expression values for both the *5p* and *3p* arms of annotated miRNA hairpins, together with metadata fields specifying Biotype, Organ_system, and Tissue. For every miRNA-tissue pair, the total abundance was computed as the sum of *5p* and *3p* expression values. To reduce noise from low-coverage observations, entries with total expression below the 25th percentile of the global abundance distribution were excluded prior to classification. Within each tissue, the relative contribution of the *5p* and *3p* arms was determined as:

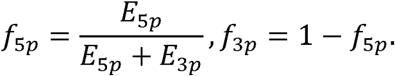

A miRNA was designated as: *5p-Singleton* if f_Sp_ 2: 0.90 and f_3p_ < 0.90, *3p-Singleton* if f_3p_ 2: 0.90 and f_Sp_ < 0.90, and *Switcher* otherwise. A 90% strand-dominance threshold was applied uniformly to all tissues. Each tissue-specific matrix was converted into a long-format table listing the dominant strand for every miRNA across organs and cell types. A global classification for each miRNA was then obtained by majority vote among all tissues and cell types that met the expression threshold. Concordance between our ML predictions and atlas-derived classifications was assessed across all miRNAs classified as *5p-* or *3p-Singletons* (excluding *Switchers*). Overall accuracy was computed as:

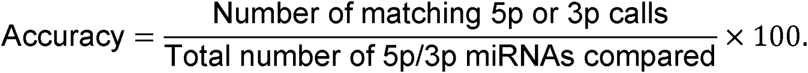

At the 90% atlas cutoff, the observed global agreement between experimental and ML model-predicted strand identity was 68.81% (median). All analyses were conducted in Python 3.10 using pandas 2.2, numpy 1.26, and matplotlib 3.9 within a Google Colab environment.

## Results

### A novel High-Throughput (HT) qPCR approach to study miRNA biogenesis

Identifying miRNAs in somatic tissues is challenging due to their small size, low abundance, and short half-life. To overcome this, we developed a high-throughput qPCR method that reliably quantifies miRNA expression and distinguishes 5p from 3p strand usage. This method is a high-throughput adaptation and general improvement of an existing methodology (26), which uses the polyuridine polymerase enzyme to elongate a polyU tail to all RNA transcripts in a given sample. This step extends the length of each miRNA present in the sample, which is then used as a template in reverse transcriptase reactions using a specially engineered universal stem-loop reverse primer (SL-polyA) (**Main Fig. 1A**).

**Main Figure 1:**
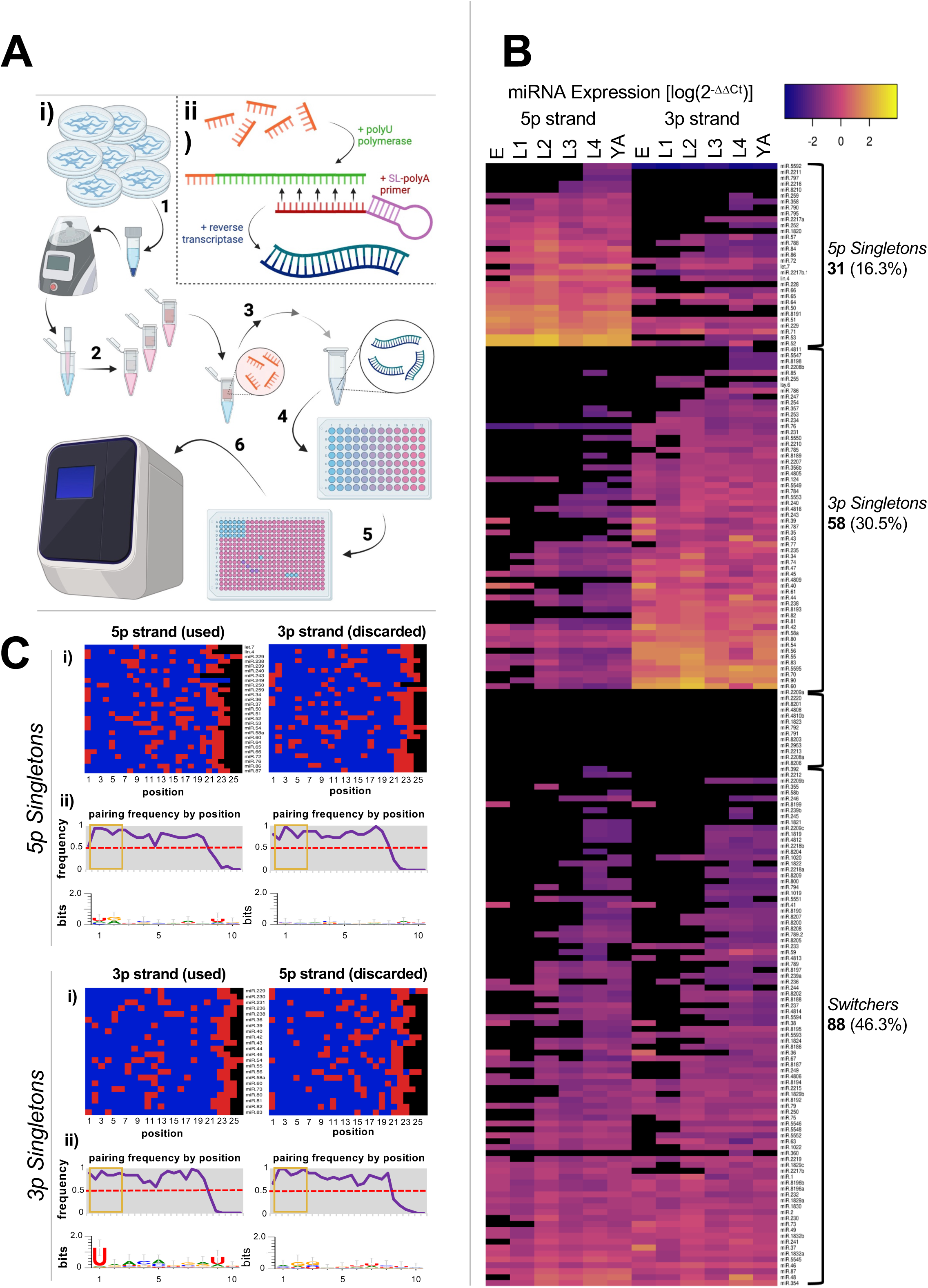
HiTmiSS Assay and Analysis on Mixed Tissue. **(A)** HiTmiSS protocol for RNA extraction, extension, reverse transcription, and quantification in high throughput (i). Steps include (1) *C. elegans* growth and homogenization, (2) RNA extraction and isolation, (3) RNA polyuridinylation, extension, and reverse transcription (details shown in ii; orange: miRNAs, green: polyU, purple: SL-polyA reverse universal primer), (4) addition of 5p or 3p miRNA-strand-specific forward primers, (5) loading into 384-well plates, and (6) quantification. **(B)** Heatmap of miRNA expression (n=3) in mixed tissue clustered by quantity, strand use, and developmental stage. Stages include embryo (E), larval 1 through 4 (L1-L4), and young adult (YA). The heatmap legend represents the log_10_ of expression values for each miRNA strand, normalized by the spike-in GFP control, at each stage, ranging from −4 (deep purple) to 4 (bright yellow). Black color indicates instances of data that fall below the strict expression threshold (one standard deviation above the mean of all expression values at that stage). **(C)** Heatmap of pairing frequency by position for the used (left) and discarded (right) strand of the 20 most highly quantified *5p* and *3p Singletons* in our mixed tissue dataset. (i) blue: paired; red: unpaired). (ii) Average pairing frequency, and logo plot of nucleotide frequency at each position displayed below (gold box marks the seed regions).

We named this method <u>Hi</u>gh-<u>T</u>hroughput <u>mi</u>RNA <u>S</u>trand <u>S</u>election (HiTmiSS) assay. HiTmiSS assay enables rapid, precise quantification of the full miRNAome and accurately distinguishes 5p from 3p strand expression (**Main Fig. 1**).

We employed this assay to systematically quantify miRNA occurrence and the usage of 5p and 3p strands in known miRNAs throughout *C. elegans* development. We extracted total RNA from *C. elegans* at six developmental stages (embryos, larval stages 1-4, and young adults) and analyzed the miRNA transcriptome and strand selection for 190 miRNAs, representing 75% of all miRNAs annotated in miRbase (25), using 380 unique forward primers across four plates (**Supplemental Table S1**).

We performed 7,980 qPCR experiments independently, querying these 190 miRNAs throughout development in triplicate (1,140 experiments at each stage) (**Main Fig. 1B, Supplemental Fig. S1**). Using stringent filters (see Materials and Methods), we detected the presence of 177 of these miRNAs (93.2%) in at least one developmental stage (**Supplemental Table S2**). The most significant number of miRNAs occurred during the L4 stage (81%), followed by young adult (75.5%), L3 (65%), L2 (56.8%), and lastly, embryo and L1 (41.6% each) stages. This aligned with previous literature suggesting that many *C. elegans* miRNAs are tissue-specific, only becoming more abundant upon the development of those tissues later in the developmental cycle (35).

As previously reported, miRNAs such as the key regulators of embryonic lethality, *miR-35* and *miR-51* families, were most abundant during embryonic and early larval stages (36). In contrast, *let-7* and *lin-4*, which control adult phenotypes, peaked at the L4 and young adult stages (37). Notably, 13 miRNAs, including *miR-791*, *miR-792*, and *miR-8206*, showed no significant expression, likely due to low abundance or being mistakenly predicted to exist.

### HiTmiSS Enhances miRNA Resolution

To benchmark the performance of HiTmiSS, we compared it to Illumina RNA-seq datasets from Kato et al. (30) and Stubna et al. (31), which profile miRNA expression across all *C. elegans* developmental stages (**Supplemental Fig. S2-S4**). HiTmiSS captured 88% of the miRNA strands identified by Kato et al., and 87% identified by Stubna et al., (overlap with both datasets = 81.6%), demonstrating strong overall concordance (**Supplemental Fig. S2A)**. Importantly, HiTmiSS more clearly resolved the top-expressed miRNAs, particularly during early development, and provided sharper 5p/3p expression profiles across stages (**Supplemental Fig. S2B**). These improvements likely stem from the usage of strand-specific qPCR primers and a standardized workflow that minimizes ligation and mapping biases. Kato et al., and Stubna et al., display broader contrasts across developmental stages (**Supplemental Fig. S2C**), reflecting the wider dynamic range that is typical of sequencing-based quantification. In comparison, HiTmiSS values are more compressed, but they reveal smoother and more gradual transitions between adjacent stages, which is characteristic of targeted qPCR-based measurements. Despite these platform-related differences, the stage-to-stage expression patterns are highly consistent across all three datasets. Many embryonic miRNAs transition from low (blue) to high (red) levels as worms progress through larval development, while a distinct subset instead peaks in young adults (YA). The distributions of values overlap almost perfectly after normalization, and the developmental trajectories of canonical miRNAs (*let-7*, *lin-4*, *miR-35* cluster, etc.) match very closely between all three datasets (**Supplemental Fig. S3**). Density plots show distinct global expression distributions for each dataset (**Supplemental Fig. S4**). Stubna et al. is enriched for low-expressed miRNAs, Kato et al. displays a tighter peak of moderate expression, and HiTmiSS covers the widest dynamic range with extended low and high tails, reflecting methodological differences between RNA-seq and targeted HiTmiSS quantification (**Supplemental Fig. S4A**). Boxplots analysis indicates comparable medians across platforms, but HiTmiSS exhibits a broader interquartile range and whiskers, consistent with increased sensitivity toward both low- and high-abundance strands relative to RNA-seq (**Supplemental Figs. S4B-C**). Importantly, most HiTmiSS-defined *5p-* and *3p-Singletons* are reproduced in Stubna et al. (73-88% of 5p and 62-79% of 3p), and developmental profiles of representative miRNAs (e.g., *let-7*, *lin-4*, *miR-51*, *miR-42*, *lys-6*, *miR-82*) illustrate clear strand dominance and stage-specific dynamics, highlighting the resolution of the HiTmiSS platform (**Supplemental Fig. S4D**).

Together, these results show that HiTmiSS faithfully recapitulates the developmental miRNA landscape captured by RNA-seq, but with enhanced clarity at the strand and stage level. Rather than serving as a redundant measurement, HiTmiSS adds resolution: it sharpens 5p/3p distinctions, smooths developmental transitions, and identifies strand dominance in agreement with independent datasets. Thus, RNA-seq provides breadth, but HiTmiSS delivers precision, making it an ideal platform for quantitative, strand-specific miRNA profiling in defined biological contexts.

### Many miRNAs show consistent strand preference throughout development

Many miRNAs identified consistently use either the 5p or 3p strand as the guide strand, which we termed ‘*Singletons*’, while others switched between 5p and 3p usage across different developmental stages. We named the second category ‘*Switchers*’.

The screen produced hits for 177 miRNAs, which we divided into *5p Singletons*, *3p Singletons*, or *Switchers* (**Main Fig. 1B and Supplemental Table S2**).

The largest group identified was miRNA *Switchers* (46.3%, 88 hits). These include members of the *miR-2* family and other regulators of developmental timing and stress responses (37,38). The remaining hits were *Singletons*, with a notable bias: *3p Singletons* (58 hits, 30.5%) outnumbered *5p Singletons* by nearly twofold. These include *miR-35* and *miR-51* (required for embryonic development), *miR-44* (stress/starvation), and *miR-58* (locomotion and dauer entry) (36,37,39). Several abundant *3p Singletons*, including *miR-60*, *miR-70*, and *miR-90*, remain functionally uncharacterized but are linked to stress response (30,38,40).

*5p Singletons* were the least common group, with 31 hits (16.3%), including the well-studied families *let-7* and *lin-4*, which are key regulators of developmental timing (**Main Fig. 1B**). Also detected were *miR-51* (embryonic lethality) and *miR-64* (heat shock response) (36,40). While some families (*let-7* and *miR-64*) consistently use the same strand, others span multiple categories. For instance, members of the *miR-51* seed family (*miR-51* to *miR-56*) and *lsy-6* exhibited consistent strand usage, with *miR-51 to miR-53* using the 5p strand and *miR-54* to *miR-56* and *lsy-6* using the 3p strand, yet none showed *Switcher* behavior. Despite sharing the same seed sequence (ACCCGUA), *miR-53* and *miR-54* are processed from opposite arms and are classified into distinct families (mirGeneDB), underscoring that convergent evolution resulted in miRNAs sharing a seed despite different evolutionary origins.

Importantly, several hits produced by the HiTmiSS assay aligned with past findings on the strand preference of well-characterized miRNAs such as *let-7, lin-4* (known *5p Singletons*), and the *miR-35* family (known *3p Singletons*) (37).

To account for the strong strand preference observed among *Singletons*, we took the top 20 *5p Singletons* and the top 20 *3p Singletons*. We compared the binding signature and nucleotide composition between the strand that is used versus the strand that is discarded (**Main Fig. 1C, i and ii**). Interestingly, the pairing frequency did not vary between these miRNAs (**Main Fig. 1C, ii**), either in the 5p or 3p strand, except for the last few nucleotides at the 3’end (**Main Fig. 1C, I**, red boxes), which were unpaired in the mature duplex. When comparing the used vs. the discarded strand (**Main Fig. 1C**, left panel vs right panel), we detected no significant difference in overall pairing (p = 1.0 in *5p Singletons*, p = 0.12 in *3p Singletons*, per Welch’s t-test) for either category of *Singleton*.

Extending this analysis to all *5p Singletons*, the 5p guide strand is relatively unpaired, with an average of 4.42 mismatches across the duplex. Thus, even miRNAs that consistently select the 5p arm typically retain several internal mismatches that reduce duplex stability and favor guide loading (**Supplemental Fig. S1B**). However, the 5p strand discarded in *3p Singletons* was significantly more paired (mean = 3.31 mismatches) than the 5p strand retained in *5p Singletons* (mean = 4.32 mismatches, p = 0.01 per Welch’s t-test), indicating that 5p strands tend to be more stable (i.e., more paired) when they are not selected (**Supplemental Fig. S1B**). Conversely, the 3p strand experienced no significant difference (p=0.16) in pairing between *3p Singletons* (mean = 3.70 mismatches) and *5p Singletons* (mean = 4.32 mismatches), suggesting its stability is less critical to the strand selection decision. Taken as a whole, the duplex structures of *5p Singletons* were less paired than those of *3p Singletons*, which might suggest a divergence in regulation of these two miRNA classes during biogenesis (**Supplemental Fig. S1B**). When narrowing our analysis to just the seed region (**Main Fig. 1C**, golden boxes), while there was higher pairing than any other position, there was no difference in binding frequency between the used and discarded strands, suggesting that different rules are also in place.

Next, we compared the nucleotide composition of these seed regions, hoping to identify signatures that could explain the discrepancy in miRNA strand selection in *Singletons*. We determined the likelihood of any given nucleotide from positions 1 through 10 in each miRNA strand in the top 20 *Singletons* in each category (5p and 3p) (**Main Fig. 1C, ii**). We detected a highly conserved uracil nucleotide at position 1, as previously shown by us and others (2), though this conservation dissolves from positions 2 through 10 (**Main Fig. 1C, ii**). Importantly, this uracil nucleotide is not equally conserved between *5p Singletons* and *3p Singletons*. 80% of *3p Singletons* start with this uracil nucleotide, while less than 40% occur in *5p Singletons*. This differential presence of this uracil may be caused by a non-stochastic pre-miRNA cleavage by the *C. elegans* DCR-1, which may possess a preference to cleave at this nucleotide.

### The intestine miRNA population is small but highly interconnected

Many of the guides detected by HiTmiSS were classified as *Switchers* (∼46%). However, some of these may instead represent stage-or tissue-specific *Singletons*, meaning miRNAs that consistently use one strand within a given context, but switch strands across different contexts. In other words, these cases would still be considered *Switchers* under our definition, but they may appear as *Singletons* when examined in only one stage or tissue, potentially confounding the analysis.

To address this, we refined our approach by analyzing somatic tissues across development, focusing on the intestine, which is a large tissue with a well-characterized transcriptome (41,42).

We coupled our HiTmiSS assay with intestine-specific miRNA immunoprecipitation, focusing on the embryo vs young adult stages. More specifically, we forced the expression of the GFP-tagged *C. elegans* Argonaute ortholog *alg-1* in the intestine tissue using the intestine-specific promoter *ges-1p*, performed an α-GFP immunoprecipitation, recovered the miRNA strands loaded onto ALG-1, and used them as templates in our HiTmiSS assay (**Main Fig. 2A**). We found that of the 93 miRNAs tested, 72 (77%) were present during at least one stage within the intestine tissue (**Main Fig. 2**). Of these, 66 were *Singletons* (92%), while 6 (8%) were *Switchers* (**Supplemental Fig. S5A**).

**Main Figure 2:**
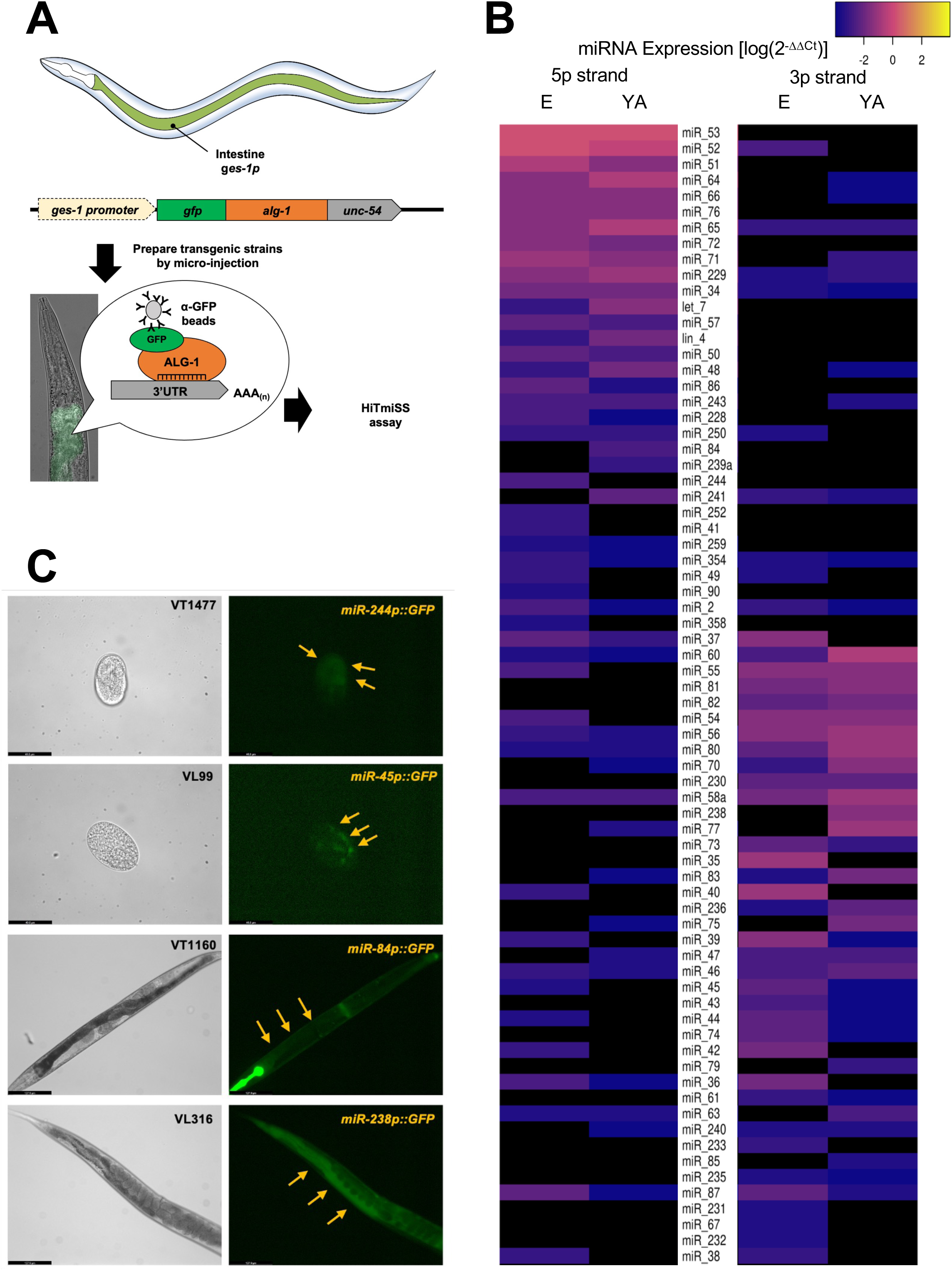
Analysis of Intestine Tissue-Specific miRNAs. **(A)** Schematics of the intestine-specific GFP-tagged ALG-1 immunoprecipitation. **(B)** Heatmap of miRNA expression in intestine tissue clustered by strand and developmental stage. The heatmap legend represents log_10_ of expression values for each miRNA strand normalized by spike-in GFP control at each stage from −4 (deep purple) to 4 (bright yellow). Black color indicates instances of data that fall below the strict expression threshold (mean of all expression values at that stage). E: embryo; YA: young adults. **(C)** Representative brightfield (left) and GFP (right) images of selected *C. elegans* strains expressing GFP under the control of promoters for miRNA known to be highly expressed in the intestine, either during the embryo stage (top two) or young adult stage (bottom two). Arrows indicate GFP expression in the intestinal tissues of both embryo and young adult stages.

This strongly differed from our whole worm data, suggesting that miRNAs previously labeled as *Switchers* may truly be *Singletons* on a tissue-by-tissue basis, only appearing as *Switchers* once all tissues are analyzed together.

Interestingly, the proportion of *5p Singletons* to *3p Singletons* was not the same in the intestinal tissue as in the mixed tissue (**Supplemental Fig. S5A**). In our initial analysis, about two-thirds of *Singletons* were *3p Singletons*, yet in the intestine tissue, *3p Singletons* were only slightly more abundant than *5p Singletons* (55% vs 45% of *Singletons* identified, respectively) (**Supplemental Fig. S5A**).

Another surprise was the presence of intestinal miRNAs with detectable expression of both 5p and 3p strands (**Main Fig. 2B**). For example, members of the *miR-51* family, well-characterized regulators of embryonic lethality (36), expressed both strands in the intestine. Similarly, *miR-44*, *miR-58*, and *miR-71* families showed dual-strand expression in intestinal samples (**Supplemental Fig. S5B**). These miRNAs appear as *Singleton* species in the total-RNA dataset, but both strands are detectable in intestine-enriched RNA, although one strand is consistently more abundant. These miRNAs therefore do not undergo strand switching; rather, they maintain dual-strand expression specifically in the intestine, potentially expanding their available 3’ UTR target space in a tissue-specific manner.

Within this dataset we also identified 6 intestine miRNAs with stage-specific strand preference; 3 *5p Singletons* (*miR-2*, *miR-87* and *miR-241*) and 3 *3p Singletons* (*miR-36*, *miR-37*, and *miR-63*) (**Supplemental Fig. S5A-B**). Importantly, four of these six *Singletons* have been independently reported in previous studies (35) (**Supplemental Fig. S5B-C**), reinforcing their biological relevance.

one life stage. These strains expressed the promoter for a single miRNA fused to GFP, and live imaging was used to validate the expression of these miRNAs at the appropriate life stage in the intestine (**Main Fig. 2C**). All four strains displayed the expected fluorescence.

### miRNA target presence does not account for strand usage

Our HiTmiSS array enables precise, strand-resolved quantification of miRNAs *in vivo.* We considered the possibility that miRNA targets could selectively stabilize one strand, thereby influencing its relative abundance and potentially confounding interpretation of the molecular basis of strand selection, rather than merely affecting HiTmiSS detection. In this model, only target-bound strands would be retained, while the unused strand would degrade, falsely suggesting strand preference.

To test this, we overexpressed *fox-1* and *hrp-2* 3’UTRs, each containing multiple binding sites for typically discarded strands (e.g., *miR-60* 5p, *miR-232* 5p, *miR-239b* 5p), in intestinal tissue (**Supplemental Fig. S6**). If target-induced stabilization were driving strand abundance, we would expect increased detection of these 5p strands. Although we observed a small increase in the abundance of some typically discarded strands (e.g. *miR-60-5p* in *fox-1* 3′UTR overexpression and *miR-232-5p* in *hrp-2* 3′UTR overexpression) we saw similar increases in canonically retained strands (e.g. *miR-60-3p* in *fox-1* overexpression), which the target stabilization hypothesis cannot explain (**Supplemental Fig. S6C**). Additionally, none of the detected changes rose above our analytical threshold of significance (i.e. no 2-fold increase or decrease) compared to wild-type controls, with 3p strands still predominating in all cases.

Taken together, these results indicate that our qPCR approach is not simply detecting stabilized miRNA strands due to the presence of targets. Instead, the relative strand abundance measured by the HiTmiSS assay likely reflects genuine strand selection, rather than artifacts introduced by target availability.

### 5p *Singletons* and *3p Singletons* follow different rules for selecting strand usage

Next, we examined the structural features associated with *Singletons*, which may explain the strict and consistent strand usage observed in these miRNAs. When comparing the used and the discarded strand, the pairing frequency varied significantly within each *Singleton*, revealing new conserved patterns (**Main Fig. 3**). Specifically, the strand used in *Singletons* often had an unpaired nucleotide at position 12, while the discarded strand did not (**Main Fig. 3A**, yellow boxes). This lack of pairing in position 12 in both *5p Singletons* and *3p Singletons* usage is intriguing. In humans, specific Argonaute proteins perform cleavage of target RNAs between positions 10 and 11, which corresponds to positions 10 and 11 along the bound miRNA strand by Watson-Crick pairing (43). *C. elegans* ALG-1 is largely homologous to human Argonaute proteins, and it may retain this cleavage activity at similar target positions, leading to the degradation of the unused strand due to its Watson-Crick base pairing at this crucial position. In addition, we detected differences in the binding frequency of the first-position nucleotide in both *5p Singletons* and *3p Singletons* (**Main Fig. 3A**, blue boxes). *3p Singletons* also exhibited unique pairings at positions 13-19 (**Main Fig. 3**, white box) with a higher pairing frequency than *5p Singletons* at the same position. These patterns are most clearly seen in specific *Singleton* miRNAs, both *5p Singletons* such as *miR-52* and *3p Singletons* such as *miR-75* (**Main Fig. 3B**).

**Main Figure 3:**
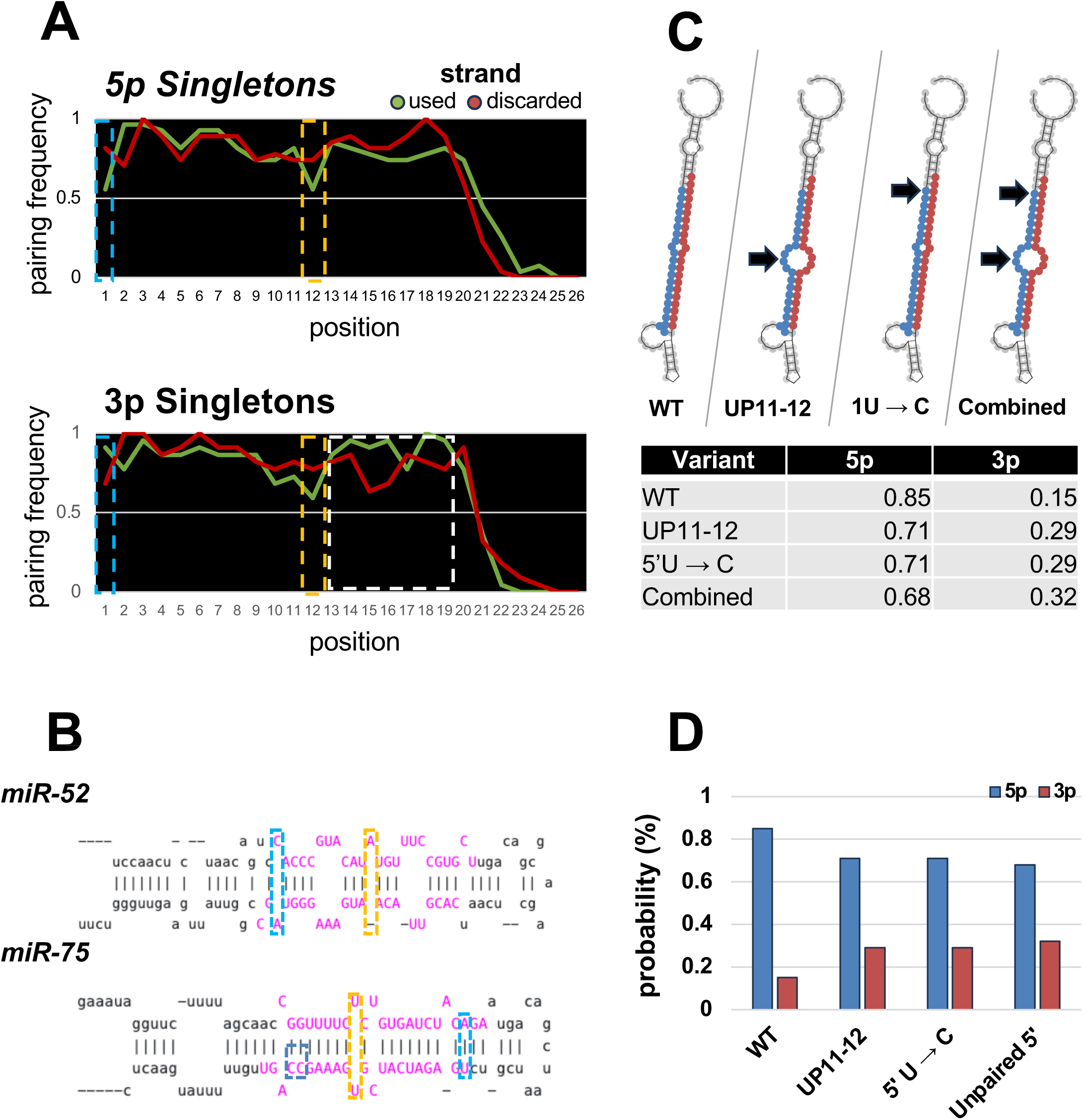
Structural and Thermodynamic Determinants of Strand Selection. **(A)** Pairing frequency by position of the used (green) and discarded (red) strand for the most highly quantified *5p Singletons* and *3p Singletons* in mixed tissue (n=20). Positions at which pairing frequency most differs are boxed, including the first nucleotide (light blue), the center of the duplex (gold), and the end of the strand (white). **(B)** Representative pri-miRNA hairpins for a *5p Singleton* (*miR-52*) and a *3p Singleton* (*miR-75*) demonstrating the occurrence of trends in pairing as described above. Mature duplex regions are highlighted in pink. **(C)** Changes in *Singleton* predictive confidence for the miRNA *let-7* as a result of *in silico* site-directed mutations. RNAfold secondary structures of pri-miRNA with mature 5p strand (blue) and mature 3p strand (red) from left to right: wildtype (WT), unpaired nucleotides at positions 11 and 12 via GG→UU substitution (UP11-12), first nucleotide U→C substitution (1U → C), combination of both prior mutations (Combined). The table summarizes the predicted strand preference by our ML model for each mutation. **(D)** Bar chart displaying probability as described above.

### An updated machine learning (ML) model for miRNA strand selection

We wanted to extract patterns and features that make *Singletons* so unique and predispose them to always use one strand over the other. To achieve this, we utilized our experimental *Singleton* data as a training set for a machine learning model designed to predict 5p versus 3p strand usage in miRNA hairpins. We hypothesized that by training such a model, we could identify the unique intrinsic features that characterize *Singletons*.

We opted for a Random Forest classifier, which we trained using our experimentally validated *Singletons* identified through our HITmiSS assay. This model incorporated 77 features, systematically organized into five functional categories (dinucleotide composition, first/last nucleotide bias, GC content and entropy, base pairing properties, and general structural features) that together capture critical aspects of miRNA mature duplex structure (**Supplemental Fig. S7–S9, Supplemental Tables S5-S6, and Materials and Methods**). The model demonstrated strong performance in distinguishing 5p from 3p usage across a set of known miRNAs and other simpler models (**Supplemental Fig. S8 and Materials and Methods**).

### 5p and *3p Singletons* possess distinct structural features

Our ML model, built using our *Singleton* data, identified several intrinsic features in these miRNAs. To generalize this model across all *C. elegans* miRNAs and study how *Switchers* can bypass these rules to switch miRNA strand preference, we applied our ML model and conducted a comparative feature analysis across the entire *C. elegans* miRNAome (374 miRNAs, miRbase).

We extracted and calculated all 77 features used in our ML model for these miRNAs (**Supplemental Table S3**), maintaining a stringent definition of *Singletons* to enhance the signal (confidence score > 80%).

As expected, we identified 54 *5p Singletons*, 52 *3p Singletons,* and 268 *Switchers* (**Main Figs. 4A-B** and **Supplemental Table S2**). Out of 77 extracted structural and sequence features, 13 exhibited statistically significant differences between the 5p and 3p strands (p < 0.05, Welch’s t-test) (**Main Figs. 4B-E**). The most pronounced difference was observed in the identity of the first nucleotide of the 3p strand, which in *3p Singletons* was consistently a uracil (p = 3.98 × 10^−^□), followed by an enrichment of cytosine in the last position (p = 5.69 × 10^−^□) (**Main Figs. 4C-4E**). Additionally, 3p strands showed an enrichment for specific motifs, particularly GAG (p = 2.51 × 10^−^□) (**Main Fig. 4E**). We also observed compositional differences: 3p strands displayed increased levels of cytosine at the first position (not shown) and tended to be slightly shorter than their 5p counterparts (p = 8.90 × 10⁻³) (**Main Fig. 4D-E**).

**Main Figure 4:**
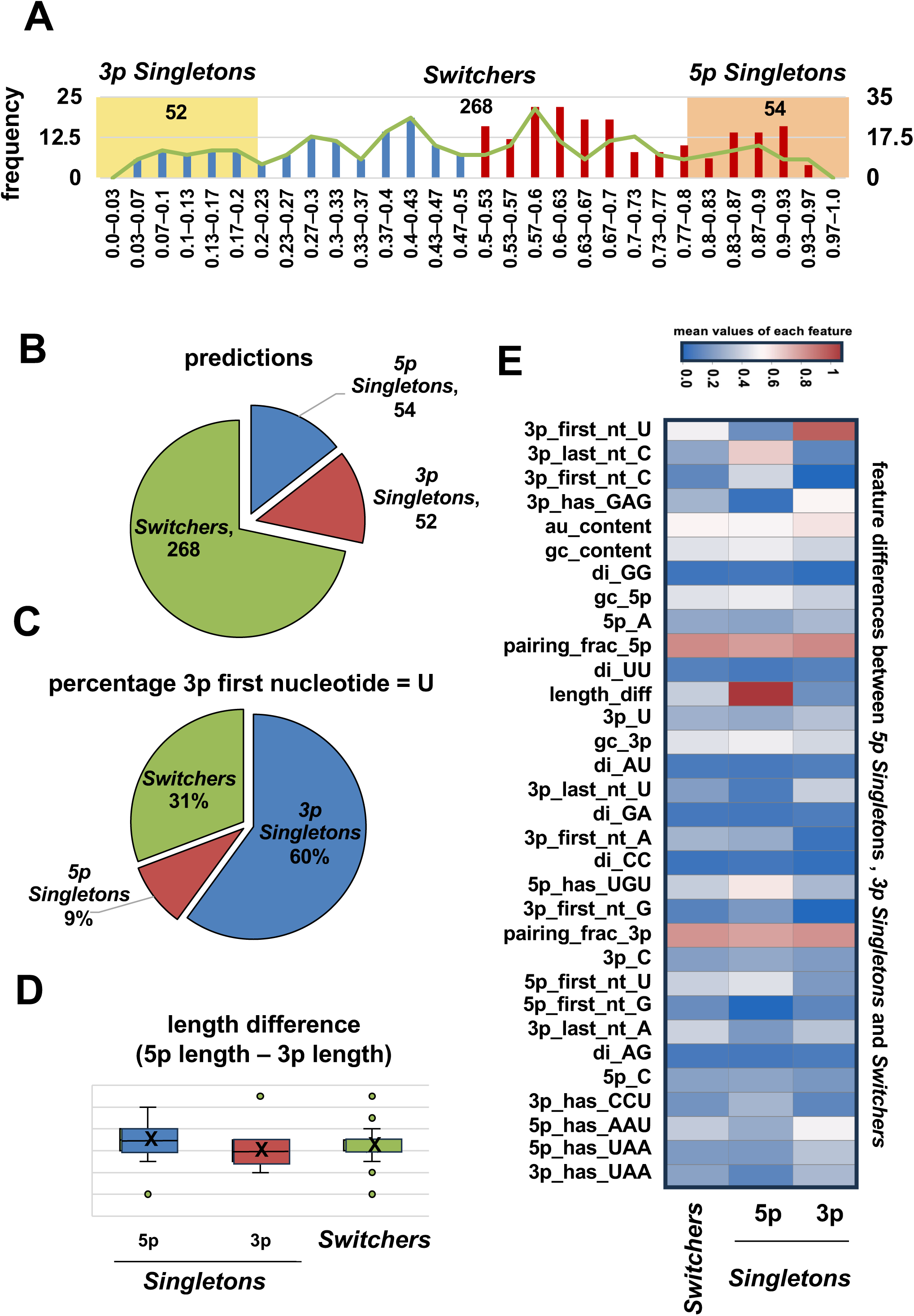
Analysis of Mature miRNA Duplex Features in *C. elegans*. **(A)** Frequency distribution of *5p Singleton* predicted confidence by our ML model across all miRNAs in *C. elegans*. *3p Singletons* (yellow) are defined as miRNAs falling below a predicted 5p confidence of 20%; *Switchers* are defined as miRNAs with a predicted 5p confidence between 20% and 80%; *5p Singletons* (red) are defined as miRNAs with a predicted 5p confidence of 80% or greater. **(B)** Predicted categorization of all *C. elegans* miRNAs by our ML model as per the above threshold. **(C)** Predicted categorization of *C. elegans* miRNAs for which uracil is the first nucleotide of the 3p strand. **(D)** Length difference (5p strand - 3p strand) for each category of *C. elegans* miRNAs. *5p Singletons* display significantly (p<0.05) longer 5p strands than 3p strands when compared either to *3p Singletons* or to *Switchers*. **(E)** Heatmap of 32 features including GC content, miRNA length, hydrogen bonding, nucleotide pairing, strand stability, and MFOLD-derived parameters enrichment in *5p* and *3p Singletons*. Heatmap values represent the mean values of each feature across the miRNA categories. For binary features (i.e., 5p_has_AAU, which is either 0 or 1), the number represents the proportion (e.g., 0.3 = 30% of miRNAs in that category have an AAU motif in the 5p strand). For continuous features (i.e., gc_content, pairing_frac_5p), the number is the average value across all miRNAs in that category. For nucleotide composition features (i.e., 5p_A, 3p_U), the number reflects the fraction of that nucleotide within the strand, features clustered by average linkage.

Surprisingly, our ML model did not cluster all the *Switchers* around a confidence score of ∼50%. Instead, we see a multi-modal distribution with several periodical clusters (**Main Fig. 4A**). This suggests that across this category, structural and thermodynamic features alone cannot fully explain strand usage, and extrinsic factors could move the needle between 5p or 3p choice in these miRNAs, perhaps depending on tissue or developmental context.

We next wanted to study the intrinsic features identified by our ML model. For these experiments, we used *Singletons* to test if altering these rules could convert them into *Switchers*. In *let-7*, modifying nucleotide pairing at positions 11–12 (UP11-12) or changing the first base (5’U→C) reduced its predicted *5p Singleton* confidence by up to 17%, consistent with our model (**Main Figs. 3C-D**). Similar results were observed in *lin-4, miR-52*, *miR-75*, and *miR-35*: introducing bulges near the center was predicted to disrupt strand preference, while removing them was predicted to reinforce it (**Supplemental Fig. S9**), highlighting the central region’s role in strand selection.

The first nucleotide’s identity and pairing also influenced strand prediction, though with variable impact across miRNAs (**Supplemental Fig. S9**). Mutating this position was predicted to alter strand usage by less than 10% in *5p Single*tons but up to 30% in *3p Singletons*, mirroring HiTmiSS results.

### Strand usage bias differs by species

Our ML model identified several features that may distinguish *Singletons* from *Switchers* in *C. elegans*. Since many miRNA sequences are conserved across species, we hypothesized that the features identified by our ML model were selectively maintained throughout evolutionary time. To do that, we applied the same model to the complete human miRNAome (**Main Fig. 5**) (**Supplemental Table S6**), and to other species throughout the tree of animal life (*Drosophila melanogaster*; fly, *Danio rerio*; zebrafish, *Xenopus laevis*; frog, *Rattus norvegicus*; rat, *Mus musculus*; mouse) (**Main Fig. 5, Supplemental Fig. S10** and **Supplemental Table S6**). While miRNA strand usage in worm, fly, zebrafish, and frog was predicted to remain relatively balanced between the 5p and 3p arms, mammalian species were predicted to exhibit a pronounced preference for the 5p strand **(Supplemental Fig. S10A)**. In *C. elegans*, only 51.3% of miRNAs were predicted to use the 5p strand, and in *Drosophila*, 55.6% were 5p. In contrast, this proportion rose sharply to 78.2% in mice and 75.9% in humans, indicating a shift toward a fixed strand preference (**Main Fig. 5A** and **Supplemental Fig. S10A)**.

**Main Figure 5:**
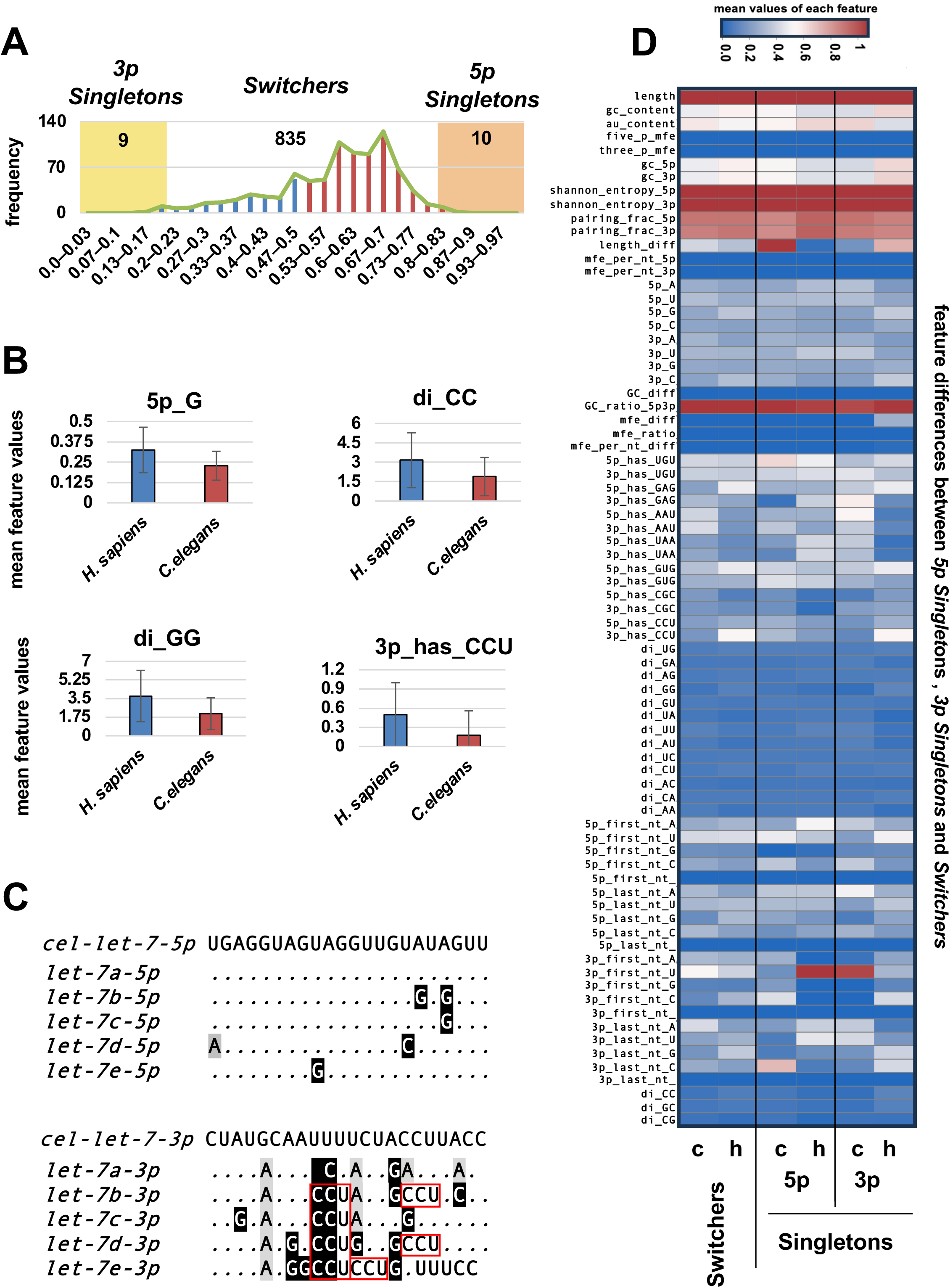
Comparative Analysis of Mature miRNA Duplex Features between *C. elegans* and *H. sapiens*. **(A)** Frequency distribution of predicted *5p Singleton* confidence across all miRNAs in *H. sapiens*. *3p Singletons* (yellow) are defined as miRNAs falling below a predicted 5p confidence of 20% or below; *Switchers* are defined as miRNAs with a predicted 5p confidence between 20% and 80%; *5p Singletons* (red) are defined as miRNAs with a predicted 5p confidence of 80% or greater. **(B)** The four most significant (p<0.05) structural differences between *H. sapiens* (left) and *C. elegans* (right) mature miRNA duplexes. From top left to bottom right: mean occurrence of guanine in the 5p strand (5p_G), mean occurrence of guanine-guanine dinucleotide in either mature strand (di_GG), mean occurrence of cytosine-cytosine dinucleotide in either mature strand (di_CC), binary classifier for presence of cytosine-cytosine-uracil trinucleotide in the 3p strand (3p_has_CCU). **(C)** Alignment of *C. elegans* (*cel*) *let-7* and five *H. sapiens* (*hsa*) *let-7* homologs shows differences in CCU occurrence within the 3p strand (red boxes). **(D)** Heatmap of differences in mean value for each of the 77 model features between each category of miRNA in *C. elegans* (c) and *H. sapiens* (h). For binary features (i.e., 3p_has_CCU, which is either 0 or 1), the number represents the proportion (e.g., 0.3 = 30% of miRNAs in that category have a CCU motif in the 3p strand). For continuous features (i.e., gc_content, au_content), the number is the average value across all miRNAs in that category. For nucleotide composition features (i.e., 5p_A, 5p_U), the number reflects the fraction of that nucleotide within the strand. Features clustered by average linkage.

Across species, we also observed an apparent decrease in pre-miRNA length of the used strands, especially in *Switchers* and *3p Singletons* (**Supplemental Fig. S10B**). This suggests that in vertebrates, shorter pre-miRNAs can adopt fewer folding patterns, potentially enhancing the use of specific strands.

Next, we sought to identify structural features that could explain the sharp increase in predicted 5p strand usage observed in mammals. We identified 13 significantly different features between worms and humans (p < 0.05, Welch’s t-test), with striking enrichment of guanine content (p = 5.62 × 10⁻²) and GG dinucleotides (p = 1.94 × 10^-2^□) on human 5p strands (**Main Fig. 5B**). Invertebrates exhibited a subtle enrichment in AU-rich motifs (UU: +2.46% higher in invertebrates; AU: +2.39% UA: +1.70%; AA: +2.11%), consistent with lower duplex stability and more flexible strand usage (**Supplemental Fig. S10C-D**). In contrast, mammals showed increased frequencies of GC-rich dinucleotides such as GG and GC, especially within the disfavored strand (di_GG: 3.6% higher in mammals than in invertebrates; di_GC: 0.9% higher in mammals) (**Main Fig. 5B**, and **Supplemental Fig. S10C-D**). The overall GC content in worm, fly, and frog miRNAs was 44-45%, whereas mouse and human miRNAs exhibited significantly higher GC content (∼52– 53%). This overall increase in GC content, which translates to an increase in duplex stability, could be one of the most important features driving 5p strand usage in vertebrates.

Finally, analysis of 5’ nucleotide identity showed that all species retained a preference for uracil at the first nucleotide of the loaded strand (2). However, mammalian species exhibited an increase in guanine and cytosine at the first position of the 3p strand, as well as higher CCU content and diCC motifs (p < 5 × 10^−2^□). These trends are evident in conserved miRNAs, such as *let-7* (**Main Fig. 5C**), which potentially serve as a negative selection marker to suppress their incorporation into RISC.

Taken together, these features may provide structural hints as to how vertebrates adopted a fixed strand usage. Shorter and more paired duplexes may enhance the selection of *Singletons* in vertebrates, while longer and less structured sequences may allow invertebrates to increase usage of *Switchers*.

### Our ML model can accurately predict human miRNA strand selection

Our *C. elegans* ML model was also capable of identifying sequence elements and structural features in human miRNAs. To further analyze these results, we compared its predictions to expression data from the Human miRNA Tissue Atlas across 65 healthy tissues and cell lines (34) (**Main Fig. 6A-B**). Overall, the model reached a median of 68% accuracy throughout all datasets, with peaks of 81% in cell lines and 78% in organs of origin (**Main Fig. 6AC**). Most errors involved 3p-dominant miRNAs misclassified as 5p, reflecting a model bias toward 5p predictions. These misclassifications often occurred when 5p and 3p strands had similar expression levels, suggesting either model uncertainty or that these miRNAs may function as *Switchers* in humans **(Supplemental Table S6).**

**Main Figure 6:**
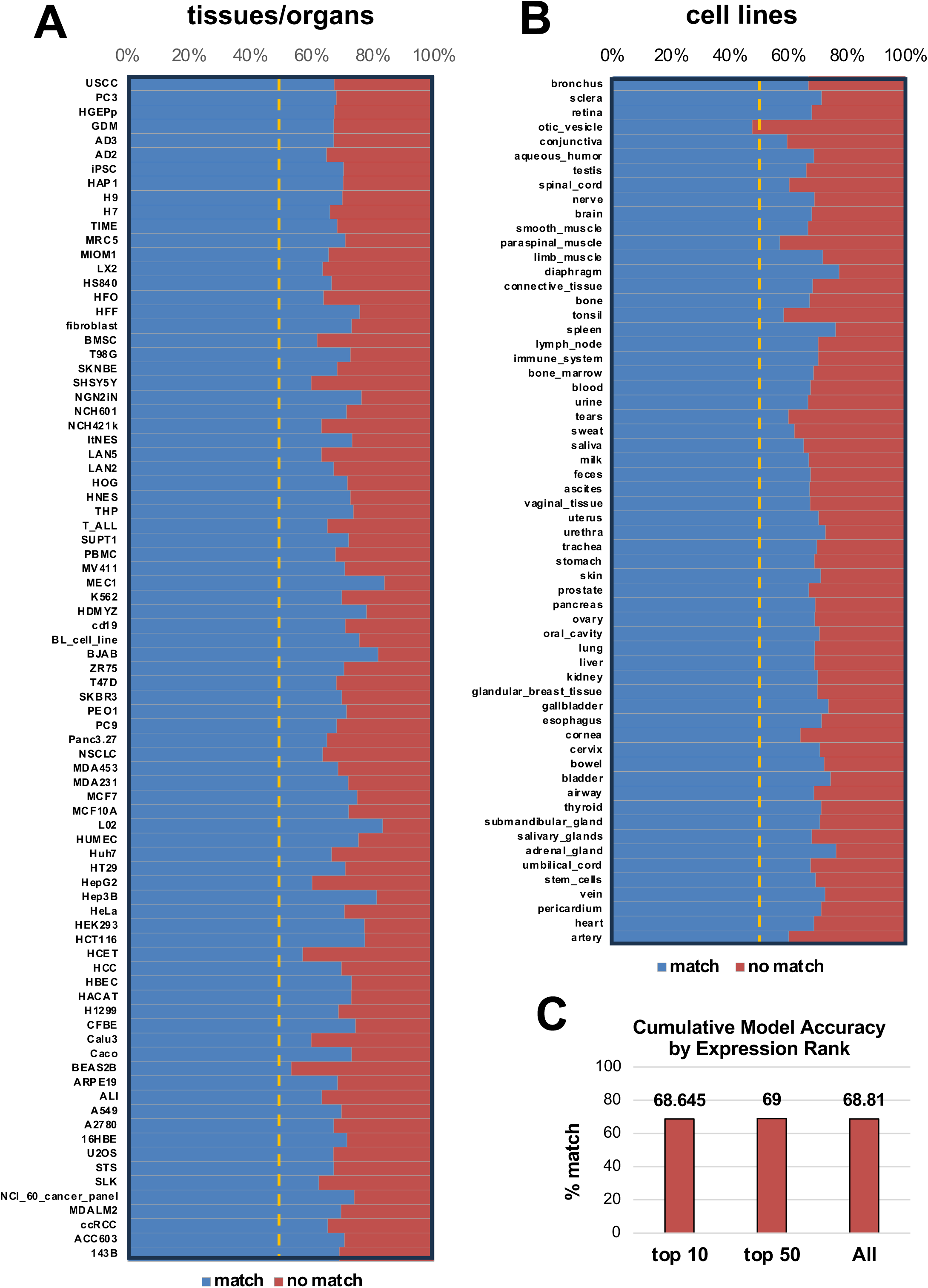
Model Accuracy for the Human miRNA Tissue Atlas Dataset. **(A)** Model accuracy expressed as a percentage of correct HiTmiSS predictions for of *H. sapiens* tissue and organ miRNAs from the miRNA Tissue Atlas (410 tissues/organs). **(B)** Model accuracy expressed as a percentage of correct HiTmiSS predictions for of *H. sapiens* cell lines miRNAs from the miRNA Tissue Atlas (264 cell lines) **(C)** Cumulative Model Accuracy by expression Rank for the top 10, 50 and all miRNAs in the miRNA Tissue Atlas.

Notably, two misclassified miRNAs, *miR-182* and *miR-378a*, are known *Switchers in vivo* (44,45). In other cases, mispredictions occurred when the model’s confidence scores for 5p and 3p usage were nearly equal (±10%). For instance, *miR-199a* and *miR-199b*, involved in hypoxia and cardiac regulation (46), were predicted as 5p dominant despite showing 3p expression and a narrow confidence margin (∼7%). Similarly, *miR-124*, critical for neurogenesis, was misclassified with a 6% confidence gap, despite clear 3p dominance (47).

Still, many ML model predicted miRNA strands were accurately detected across diverse human tissues, including key regulators with strong strand bias, indicating that our *C. elegans*-trained model generalizes well to vertebrates (**Main Fig. 6C and Supplemental Table S6**). For example, *miR-21*, a highly expressed oncomiR, was correctly predicted as 5p-dominant, with over 50-fold higher 5p expression than 3p (48). Similarly, *let-7a* showed a 248-fold preference for 5p and was correctly classified (49). Other highly expressed miRNAs, such as *miR-1* and miR*-141,* were also correctly predicted as 3p dominant.

Overall, these findings demonstrate that the model reliably predicts strand usage for miRNAs in healthy human tissues, particularly when strand bias is strong. Errors were enriched among duplexes with more balanced expression levels or context-dependent strand usage, highlighting the complexity of predicting which strand will be used in cases of weak or dynamic regulation of strand selection.

### Evolution drives strand-selection shifts

Because our ML model predictions in *C. elegans* mirror miRNA strand usage patterns observed in humans, we next asked whether the sequence and structural “feature rules” learned in *C. elegans* were conserved or repurposed across metazoans. Very few miRNAs are deeply conserved between nematodes and humans, and those that are (e.g., *let-7*) are uniformly *5p Singletons* in both species. A notable exception is *miR-34* (**Main Fig. 7**). In *C. elegans, miR-34* contributes to stress resistance, aging, and DNA-damage responses (50,51), while in humans it functions downstream of p53 and acts as a tumor suppressor that inhibits proliferation, prevents metastasis, and promotes apoptosis (52–54). Strikingly, *miR-34* is a *3p Singleton* in *C. elegans* across all developmental stages, yet it becomes a *5p Singleton* in mammals, matching both our ML model’s predictions and the miRNA atlas data. To understand this inversion, we traced sequence and structural changes in *miR-34* precursors across fly, mouse, and human (**Main Fig. 7**). Across metazoans, *miR-34* shows a clear evolutionary flip from 3p maturation in basal organisms to 5p dominance in mammals. In *C. elegans*, *miR-34* is predominantly processed from the 3p arm (P(3p)=0.76), driven by a strong thermodynamic bias. The 3p arm contains 63.6% GC versus 54.5% on the 5p arm, yielding a more stable 3′ duplex end and favoring Argonaute loading (**Main Fig. 7**). Although the 5p product begins with a favorable 5′-A, duplex asymmetry still enforces 3p selection. A decisive shift appears in insects: *Drosophila miR-34-5p* acquires a 5′-U, which is the canonical AGO-loading nucleotide, coincident with a switch to 5p dominance (P(5p)=0.68). This A-to-U substitution at position 1 is retained in vertebrates, stabilizing 5p usage (mouse P(5p)=0.70; human miR-34a frequently ≥0.7) (**Main Fig. 7**). In addition, human precursors display shorter terminal loops (∼6–8 nt versus ∼12–15 nt in worm), promoting precise Drosha/Dicer cleavage and a perfectly conserved 5′ seed start. Dinucleotide composition also evolves in a direction that weakens 5p end-pairing: di_GU progressively increases, introducing GU wobble near the guide and lowering local duplex stability, which favors 5p loading. Conversely, di_CG and di_GA decline to zero in mammals (di_GA is already lost in fly), eliminating stabilizing stacking interactions still present in worms. Collectively, the shift from 3p to 5p usage in *miR-34* occurred through (i) loss of 3p GC enrichment, (ii) gain of AU-rich 5′ nucleotides that enhance AGO recognition, (iii) terminal loop shortening that improves processing precision, and (iv) dinucleotide substitutions that reduce 5p duplex stability relative to 3p. Thus, fly represents an intermediate state, while mouse and human complete the evolutionary inversion, establishing *miR-34a-5p* as the conserved, functionally dominant product in mammals (**Main Fig. 7**).

**Main Figure 7:**
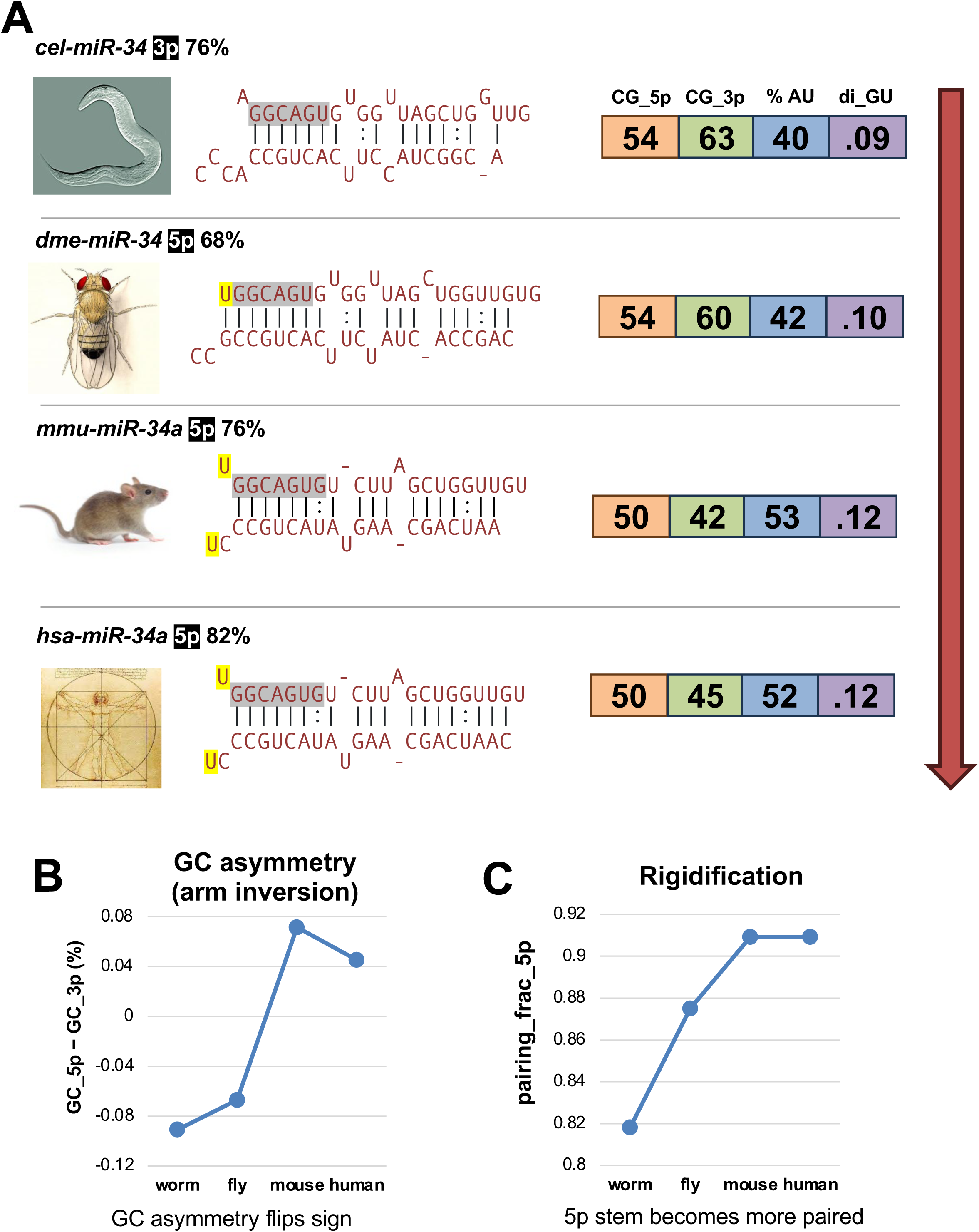
Evolutionary analysis of *miR-34* strand usage in worms, flies, mice, and humans. Across metazoans, *miR-34* undergoes a striking evolutionary inversion from *3p Singleton* in *C. elegans* to *5p Singleton* in insects and mammals. (Top) The seed region of *miR-34* is highlighted in gray. In worms, stronger 3p GC content (63.6% vs. 54.5% on 5p) enforces 3p loading despite a 5′-A on the 5p guide. In insects, *miR-34-5p* acquires a 5′-U (yellow), coincident with 5p dominance (P(5p)=0.68), and this feature is conserved in vertebrates (mouse P(5p)=0.70; human ≥0.7. (Bottom Left). Model-derived GC asymmetry rises gradually across species, from *C. elegans* to fly, mouse, and human, and this trend is mirrored by a progressive increase in 5p pairing (bottom right).

### A community resource to study miRNA strand selection

To support the broader research community, we have made all components of our strand selection framework openly available via GitHub (https://github.com/MangoneLab/Caligula_app). This resource includes our trained ML model, as well as Python scripts for both local, cumulative, and browser-based use. The platform is designed for flexibility and ease of use, allowing users to input a single miRNA or a batch of sequences in FASTA format to obtain strand selection predictions. We also provide pre-processed strand selection predictions, including all 77 features used by our ML model, for every miRNA annotated in miRBase across seven species: worm, fly, zebrafish, frog, mouse, rat, and human (**Supplemental Tables S3 and S6**).

By making these tools and datasets accessible, we aim to provide a practical, scalable, and user-friendly solution for investigating miRNA strand usage across systems and species. This represents the first publicly available platform for high-throughput, strand-specific miRNA analysis, enabling researchers to integrate strand selection into both basic and translational studies.

## Discussion

This work establishes a unified, experimentally validated model of miRNA strand selection that links intrinsic duplex structure to context-dependent strand usage across development and tissues. By combining ∼8,000 qPCR measurements with feature-based machine learning, we show that strand choice is highly regulated and not stochastic, and that conserved sequence signatures help enforce *Singleton* behavior while *Switchers* remain responsive to developmental and tissue cues.

Compared to RNA-seq, HiTmiSS offers distinct advantages for miRNA analysis (**Supplemental Figs. S4-S6**). The primary differences between these methods lie in the output produced. In RNA-seq, the output consists of sequences that must be mapped onto a reference genome to infer strand usage and abundance, whereas in HiTmiSS, the output provides direct quantification of each strand. Because strand-specific primers directly amplify each mature duplex arm, HiTmiSS avoids inference-based strand assignment and ligation bias, which can obscure low-abundance species or inflate the apparent abundance of highly expressed miRNA families. We find that HiTmiSS is highly sensitive and reproducible, detecting transcripts that are often weakly captured by RNA-seq while still recapitulating developmental trends observed in sequencing datasets. Thus, RNA-seq provides breadth and discovery power, whereas HiTmiSS delivers quantitative precision and clearer 5p/3p discrimination, especially in low-abundance or tissue-restricted contexts.

Recent approaches, such as AQ-seq, have improved small RNA-seq protocols by reducing ligation bias and enabling isomiR detection (55). However, because AQ-seq still relies on sequencing-based inference for strand assignment, it may be affected by residual mapping ambiguities. In contrast, HiTmiSS directly quantifies 5p and 3p strands using strand-specific primers, offering a more targeted approach to strand-resolved analysis. Its combination of sensitivity, quantitative precision, and reproducibility makes it well-suited for studying miRNA strand selection across tissues and developmental stages.

Our study enabled the classification of miRNA into *5p Singletons*, *3p Singletons*, or *Switchers*, informing a predictive model validated against human miRNA expression data.

We found that in *C. elegans,* half of miRNAs (46%) behave as *Switchers*, altering strand usage across development, indicating that strand selection is context-dependent rather than fixed (**Main Fig. 1**). In contrast, key developmental regulators such as *let-7*, *lin-4*, and *miR-35* are *Singletons*, likely reflecting a need for consistent strand usage to ensure stable gene expression. Notably, *Singletons* tend to be more highly expressed (**Main Fig. 1B**), perhaps due to stronger structural or thermodynamic constraints. In the intestine, tissue-specific ALG-1 immunoprecipitation revealed a strong preference for *Singletons* (**Main Fig. 2**). This suggests that strand selection is also shaped by tissue-specific factors, such as the cellular environment or cofactors. These factors would have substantial consequences for miRNA regulatory output, as strand choice reshapes the accessible set of 3′UTR targets. While our approach captures intrinsic sequence determinants, with the caveat that only one tissue (intestine) and two developmental stages (embryo and young adult) were profiled, the tissue-specific strand usage we observe points to an additional, regulated layer of post-transcriptional control. Identifying the proteins and regulatory pathways that modulate strand choice is an important future direction, both in *C. elegans* and likely in other metazoans. Importantly, since we examined only one somatic tissue, some miRNAs classified as *Switchers* in our experiments may be tissue-specific *Singletons*.

A concern was that target mRNAs might stabilize one strand, creating an artificial strand preference in our HiTmiSS assay. To test this, we overexpressed 3’UTRs with binding sites for typically low-abundance strands in the intestine. Although we cannot rule out other downstream processes affecting miRNA stability, such as decay rates or target-mediated miRNA degradation (56,57), strand levels remained unchanged, suggesting that HiTmiSS reflects strand usage, not target-driven stabilization (**Supplemental Fig. S6**).

When comparing the structural composition of each mature duplex, a series of patterns emerged. Regarding the strand pairing, there is no universal trend showing that 5p strands were always more unpaired than 3p strands, regardless of selection. Instead, selected strands (whether 5p or 3p) tended to tolerate moderate unpairing, and unselected 5p strands (i.e., in *3p Singletons*) were more paired, suggesting that greater structural stability may discourage selection, supporting a role for pairing in strand discrimination (**Main Fig. 1C**).

*5p Singletons* tended to have the first nucleotide unpaired to its corresponding strand in the duplex (**Main Fig. 3A**). This difference in pairing repeats in the middle of the duplex and lowers toward the end (**Main Fig. 3A**). Contrarily, *3p Singletons* present a near-opposite trend, with the first nucleotide more often paired and the end portion of the sequence also paired, though the center of the sequence shows similar unpairing to *5p Singletons*.

The effects of the 5’ nucleotide identity and central region pairing on strand selection and Argonaute loading have been well known for a long time, with foundational work highlighting the importance of 5’ nucleotide identity and central region pairing for Argonaute loading (14,58,59). These studies established general rules governing strand preference, such as the thermodynamic asymmetry model, but were derived under controlled conditions, often focusing on a limited number of miRNAs in model systems. While they established the principles of strand selection, they did not provide a scalable means to measure how these rules were implemented, or overridden, in living organisms across developmental stages, tissues, or physiological contexts. HiTmiSS advances the field by shifting the focus from mechanism to measurement, offering a practical, high-throughput platform for directly quantifying strand usage across biological conditions, developmental stages, and tissues. In doing so, HiTmiSS transforms strand selection from a static rule set into a measurable, context-dependent regulatory layer.

In addition, our results added positional detail to these foundational studies by showing that asymmetry varies along the duplex. Both *5p* and *3p Singletons* show central unpairing, likely reflecting a conserved feature for Dicer or Argonaute engagement but not strand bias. Notably, *5p Singletons* show decreasing pairing from the seed to the 3’ end, while *3p Singletons* display the opposite trend. These opposing patterns suggest that specific structural cues along the duplex, not just global stability, guide strand selection, pointing to evolved, position-dependent regulatory signals within miRNA architecture.

To expand our analysis, we extracted 77 structural and compositional features from our validated *Singletons* and trained a Random Forest classifier (ML model) to predict dominant strand usage. When comparing these features (**Main Fig. 4**), we found several strand-specific differences, including a significantly higher occurrence of uracil at the first nucleotide in 3p strands and enrichment for GAG motifs.

We tested our predictions using *in silico* site-directed structural mutations in canonical miRNAs such as *let-7*, *lin-4*, and *miR-52*, *miR-75* (**Main Fig. 3C, Supplemental Fig. S9**). Disrupting features such as central bulges (**Main Fig. 3C, “UP11-12”**) or first nucleotide pairing (**Main Fig. 3C, “1U->C”**), altered predicted strand preference to some degree, further confirming their role in the selection process. Surprisingly, the degree to which they impact miRNA strand preference was approximately equal, but when disrupted in tandem, they led to an even greater loss of predicted strand preference (**Main Fig. 3C, “Combined”, Supplemental Fig. S9**). In addition, these findings phenocopy the trends observed in our experimentally validated *Singletons* (**Main Fig. 3A**). Taken together, these results suggest that the structural and thermodynamic features identified in our wet bench-validated *5p* and *3p Singletons* are indeed critical regulators of strand selection in nematodes.

It is important to note that these analyses do not constitute experimental validation of the model. Rather, they provide an additional test of the model by probing how specific sequence and structural features influence its predictions. We modified known determinants of strand usage, such as 5′ nucleotide identity, central mismatches, and local thermodynamic asymmetry, within biologically plausible ranges and consistent with feature distributions observed in the training data. We focused not on single predictions, but on systematic and reproducible shifts across multiple miRNAs and perturbation types. In several cases, including *miR-75*, the modified sequences not only reduced confidence but also flipped predicted strand preference, an outcome consistent with switcher-like behavior reported in the literature. However, we emphasize that these findings are predictive and hypothesis-generating, and experimental testing will be required to determine whether these perturbations alter strand usage *in vivo*.

Expanding this framework, we then applied our ML model to predict strand usage across the human miRNA dataset (**Main Fig. 5A**). We were particularly curious to test if the structural and thermodynamic features and motifs identified in *C. elegans* were conserved in vertebrates.

Compared to the nearly equal distribution of 5p and 3p usage in *C. elegans*, using the same cutoff of 80%, the vast majority of human miRNAs are identified as *Switchers* (831/854 predicted *Switchers*, 97.3%). However, within this class, there is a clear bias toward 5p strands, with over 75% of predictions favoring 5p strand usage (**Main Fig. 5A**). This divergence likely reflects increased specialization in miRNA maturation and Argonaute loading in vertebrates, potentially driven by evolutionary selection for strand-specific targeting or stability.

Despite the lower predictive accuracy of our ML model when applied to *H. sapiens* miRNAs (69%) compared to its performance on the *C. elegans* training set (88%) (**Main Fig. 6C**, **Supplemental Fig. S2B**), we do not interpret this as evidence of overfitting. Rather, the ability of the *C. elegans*-trained model to achieve performance substantially above random on a distantly related species suggests that key aspects of miRNA strand selection are indeed governed by conserved predictive features. While quantitative differences in model accuracy likely reflect evolutionary divergence and the influence of lineage-specific modifiers, the persistence of these core features supports the idea that a foundational set of strand selection rules is conserved. Thus, our model reveals not only species-specific differences but also a shared regulatory logic underlying miRNA strand selection that has been retained across metazoan evolution (**Supplemental Fig. S10**). Invertebrate systems appear to tolerate greater flexibility, allowing for more balanced usage of the duplex arms. In contrast, vertebrates, particularly mammals, exhibit strong sequence and structural biases that favor the selection of the 5p strand (**Supplemental Fig. S10**). This transition likely reflects the growing complexity of regulatory networks and the need for precise post-transcriptional gene control in higher organisms. As miRNA families expanded during vertebrate, particularly mammalian, evolution, selective forces likely favored stable strand usage to ensure consistent and non-redundant regulation (2). This trend would promote *Singletons* over *Switchers*, reducing regulatory variability. The fixation of 5p strand usage may thus represent stabilizing selection for predictable miRNA function, with structural features reinforcing 5p bias evolving alongside increasing organismal complexity.

In addition to this imbalance in *Singletons* vs *Switchers*, the nucleotide composition of their 5p and 3p strands is also different, with a significant enrichment of guanine content in human 5p strands and cytosine-rich motifs in 3p strands (**Main Fig. 5B**). These compositional biases are consistent with known differences in Argonaute protein specificity and may reflect lineage-specific constraints on strand functionality (60).

Importantly, these cross-species trends persisted in known miRNAs such as *let-7*, which was consistently predicted and validated as a *5p Singleton* both in *C. elegans* and in humans, although at different confidence levels. This supports the robustness of our predictive ML model and highlights conserved features of miRNA processing, even in the presence of divergent strand preferences.

A surprising trend emerges when we compare our ML model prediction in human miRNAs to the human miRNA Tissue Atlas. Our ML model developed in *C. elegans* is capable of accurately (∼70%) predicting miRNA strand usage in humans (**Main Fig. 6**). This is unexpected, considering the vast evolutionary gap (∼600 million years) between these two systems, which share less than 35% of nucleotide conservation in genes (61). Despite this, the features we have identified seem to be evolutionarily conserved.

This suggests that miRNA strand selection is, at its core, a fundamental cellular process that has evolved very little over time, even as miRNA regulatory networks expanded in complexity with the increased cellular specialization. This becomes more evident when we overlay miRNA strand usage in human tissues of different origins. Here, the same miRNA is always predicted to use the same strand in other somatic tissues (**Main Fig. 6A**, **Supplemental Table S6**). This invariance suggests that strand selection is most likely encoded by intrinsic properties of the hairpin itself, rather than by external, context-dependent cues. Such robustness implies that strand choice is not merely a byproduct of downstream regulatory needs, but a core mechanistic feature established early in miRNA biogenesis. We propose that during vertebrate evolution, particularly with the rise of tissue specialization, miRNA strand usage remained anchored to conserved structural determinants even as the broader miRNA regulatory network expanded and diversified. This mechanistic rigidity may stabilize gene regulatory outputs in the face of cellular complexity.

Tracing the origins and evolution of miRNA families remains challenging. Although miRNAs emerged early in metazoan history, multiple bursts of rapid expansion occurred during bilaterian diversification (∼520–500 Mya) and again during the vertebrate whole-genome duplications. These events generated large and highly specialized miRNA families that later diverged in sequence, genomic context, and copy number, often obscuring clear orthologous relationships between vertebrate and invertebrate miRNAs (2). The *C. elegans miR-34* family is one of the few deeply conserved examples in which such evolutionary comparisons can still be performed. Our analysis of *miR-34* (**Main Fig. 7**) shows that miRNA strand selection is not fixed but can be evolutionarily rewired through incremental sequence and structural changes to the hairpin precursor. Although the rules learned in *C. elegans* can predict strand usage in humans, the evolutionary trajectory between these species indicates that these rules are repurposed rather than strictly conserved. The transition of *miR-34* from 3p maturation in basal metazoans to 5p dominance in mammals demonstrates that relatively subtle features, including 5′ nucleotide identity, GC asymmetry, terminal loop length, and dinucleotide composition, are sufficient to invert guide-strand selection without disrupting hairpin recognition or overall function. These findings suggest that strand choice itself may be an evolvable regulatory parameter. Altering which arm generates the dominant guide effectively redefines the seed sequence and rewires target networks without deleting or duplicating the miRNA gene. Evolution can therefore modify miRNA function through “strand-switching,” expanding regulatory potential while preserving genomic economy.

The observation that intermediate species exhibit partial transitions further supports a model in which strand usage diverges gradually, as selective pressures favor a new dominant arm with distinct targets.

Importantly, the same features that govern strand selection in worms (5′-nucleotide identity, duplex asymmetry, wobble positioning, and cleavage precision) also predict usage in insects, mammals, and humans. In this view, evolution does not invent new rules; instead, it repurposes existing thermodynamic and structural logic to shift guide preference. Together, these data indicate that strand selection is a flexible, evolutionarily tunable mechanism for reshaping miRNA regulatory networks. The conservation of feature-level logic across metazoans, combined with species-specific rewiring of dominant strands, suggests that strand switching may represent a widespread but underappreciated layer of post-transcriptional innovation.

There are limitations to the present work. We profiled only one somatic tissue (the intestine) and two developmental stages, and it is likely that additional *Singletons* remain hidden within other tissues or stress environments. The ML model currently captures intrinsic features but not Ago preferences, cofactors, RNA modifications, or degradation pathways that may bias strand retention. Identifying these regulators, and defining how they interface with duplex structure, will be an important future direction.

Finally, we have developed a user-friendly resource on GitHub (https://github.com/MangoneLab/Caligula_app) that allows researchers to predict miRNA strand selection using our ML model. To support broad application, we also provide pre-processed strand selection predictions with all 77 input features for all miRNAs in miRBase across seven species, including worm, fly, zebrafish, frog, mouse, rat, and human (**Supplemental Tables S3 and S6**).

In summary, this study establishes a predictive and experimentally validated framework for miRNA strand selection, combining high-throughput qPCR (HiTmiSS) with an ML model to uncover both conserved and context-dependent rules governing strand usage. Unlike previous studies that inferred strand selection indirectly or focused on a limited set of miRNAs, our study provided strand-specific quantification across developmental stages and tissues in *C. elegans*, revealing dynamic and non-stochastic strand usage. We also produced a biologically grounded and publicly available ML model trained on curated *in vivo* strand data that accurately predicts strand selection across multiple species, including vertebrates. We also prepared the first species-wide probability atlas of 5p vs. 3p usage across several metazoan miRNAs, including humans, enabling genome-scale interpretation of strand bias. In addition, we highlighted a detailed dissection of 77 intrinsic features that together define a conserved, programmable regulatory logic for strand selection. This integrative approach extends beyond descriptive rules, providing a quantitative and generalizable model of strand selection applicable across metazoan evolution.

In conclusion, our findings illustrate the complexity of miRNA strand selection and reveal that this process is not governed solely by thermodynamic asymmetry. Instead, a combination of structural motifs, sequence biases, and cellular context dictates strand usage. Our work establishes a predictive framework for understanding strand selection but also opens new avenues for studying the evolution of miRNA regulatory logic across metazoans.

## Data Availability

The raw data produced by the HiTmiSS assay are included in the **Supplemental Table S2**. The ML model is included as Supplemental Data and is also available on GitHub (https://github.com/MangoneLab/Caligula_app).

## Acknowledgments

We thank Dr. Murugan for assistance in HiTmiSS assay optimization.

## Author Contributions

D.M. and M.M. designed the experiments. D.M. performed the experiments. D.M. and M.M. prepared the ML model and performed the bioinformatics analysis. H.L. supervised the development of the ML model. D.M., A.E., and H.H. performed the HiTmiSS assay experiments. D.M. and M.M. prepared the main and supplemental figures and supplemental tables. D.M. and M.M. wrote the manuscript. All authors have read and approved the manuscript.

## Supplemental Information titles and legends

***Supplemental Table S1:* Primers used in this study.**

***Supplemental Table S2:* HiTmiSS assay results.**

***Supplemental Table S3:* Features used in our ML model and their weights.**

***Supplemental Table S4:* Comparison of ML model with miRNA Tissue Atlas.**

***Supplemental Table S5: ML prediction in C. elegans.***

***Supplemental Table S6:* ML prediction in other species.**

## References

1. Bartel, D.P. (2018) Metazoan MicroRNAs. Cell, 173, 20–51.

2. Wolter, J.M., Le, H.H., Linse, A., Godlove, V.A., Nguyen, T.D., Kotagama, K., Lynch, A., Rawls, A. and Mangone, M. (2017) Evolutionary patterns of metazoan microRNAs reveal targeting principles in the let-7 and miR-10 families. Genome Res, 27, 53–63.

3. Lee, Y., Ahn, C., Han, J., Choi, H., Kim, J., Yim, J., Lee, J., Provost, P., Radmark, O., Kim, S. et al. (2003) The nuclear RNase III Drosha initiates microRNA processing. Nature, 425, 415–419.

4. Han, J., Lee, Y., Yeom, K.H., Kim, Y.K., Jin, H. and Kim, V.N. (2004) The Drosha-DGCR8 complex in primary microRNA processing. Genes Dev, 18, 3016–3027.

5. Gregory, R.I., Yan, K.P., Amuthan, G., Chendrimada, T., Doratotaj, B., Cooch, N. and Shiekhattar, R. (2004) The Microprocessor complex mediates the genesis of microRNAs. Nature, 432, 235–240.

6. Denli, A.M., Tops, B.B., Plasterk, R.H., Ketting, R.F. and Hannon, G.J. (2004) Processing of primary microRNAs by the Microprocessor complex. Nature, 432, 231–235.

7. Ladewig, E., Okamura, K., Flynt, A.S., Westholm, J.O. and Lai, E.C. (2012) Discovery of hundreds of mirtrons in mouse and human small RNA data. Genome Res, 22, 1634–1645.

8. Ruby, J.G., Jan, C.H. and Bartel, D.P. (2007) Intronic microRNA precursors that bypass Drosha processing. Nature, 448, 83–86.

9. Iwasaki, S., Kobayashi, M., Yoda, M., Sakaguchi, Y., Katsuma, S., Suzuki, T. and Tomari, Y. (2010) Hsc70/Hsp90 chaperone machinery mediates ATP-dependent RISC loading of small RNA duplexes. Mol Cell, 39, 292–299.

10. Pare, J.M., Tahbaz, N., Lopez-Orozco, J., LaPointe, P., Lasko, P. and Hobman, T.C. (2009) Hsp90 regulates the function of argonaute 2 and its recruitment to stress granules and P-bodies. Mol Biol Cell, 20, 3273–3284.

11. Nakanishi, K. (2016) Anatomy of RISC: how do small RNAs and chaperones activate Argonaute proteins? Wiley Interdiscip Rev RNA, 7, 637–660.

12. Meijer, H.A., Smith, E.M. and Bushell, M. (2014) Regulation of miRNA strand selection: follow the leader? Biochem Soc Trans, 42, 1135–1140.

13. Medley, J.C., Panzade, G. and Zinovyeva, A.Y. (2021) microRNA strand selection: Unwinding the rules. Wiley Interdiscip Rev RNA, 12, e1627.

14. Frank, F., Sonenberg, N. and Nagar, B. (2010) Structural basis for 5’-nucleotide base-specific recognition of guide RNA by human AGO2. Nature, 465, 818–822.

15. Melendez, A. and Levined, B. (August 24, 2009) In Community, T. C. e. R. (ed.), WormBook. WormBook.

16. Zinovyeva, A.Y., Veksler-Lublinsky, I., Vashisht, A.A., Wohlschlegel, J.A. and Ambros, V.R. (2015) Caenorhabditis elegans ALG-1 antimorphic mutations uncover functions for Argonaute in microRNA guide strand selection and passenger strand disposal. Proc Natl Acad Sci U S A, 112, E5271–5280.

17. Caceres, J.F., Stamm, S., Helfman, D.M. and Krainer, A.R. (1994) Regulation of alternative splicing in vivo by overexpression of antagonistic splicing factors. Science, 265, 1706–1709.

18. Mangone, M., Macmenamin, P., Zegar, C., Piano, F. and Gunsalus, K.C. (2008) UTRome.org: a platform for 3’UTR biology in C. elegans. Nucleic Acids Res, 36, D57–62.

19. Mangone, M., Manoharan, A.P., Thierry-Mieg, D., Thierry-Mieg, J., Han, T., Mackowiak, S.D., Mis, E., Zegar, C., Gutwein, M.R., Khivansara, V. et al. (2010) The landscape of C. elegans 3’UTRs. Science, 329, 432–435.

20. Steber, H.S., Gallante, C., O’Brien, S., Chiu, P.L. and Mangone, M. (2019) The C. elegans 3’ UTRome v2 resource for studying mRNA cleavage and polyadenylation, 3’-UTR biology, and miRNA targeting. Genome Res, 29, 2104–2116.

21. Murari, E., Meadows, D., Cuda, N. and Mangone, M. (2024) A comprehensive analysis of 3’UTRs in Caenorhabditis elegans. Nucleic Acids Res.

22. Kozomara, A. and Griffiths-Jones, S. (2014) miRBase: annotating high confidence microRNAs using deep sequencing data. Nucleic Acids Res, 42, D68–73.

23. Sulston, J.E., Schierenberg, E., White, J.G. and Thomson, J.N. (1983) The embryonic cell lineage of the nematode Caenorhabditis elegans. Dev Biol, 100, 64–119.

24. Frokjaer-Jensen, C., Davis, M.W., Hopkins, C.E., Newman, B.J., Thummel, J.M., Olesen, S.P., Grunnet, M. and Jorgensen, E.M. (2008) Single-copy insertion of transgenes in Caenorhabditis elegans. Nat Genet, 40, 1375–1383.

25. Kozomara, A., Birgaoanu, M. and Griffiths-Jones, S. (2019) miRBase: from microRNA sequences to function. Nucleic Acids Res, 47, D155–D162.

26. Mei, Q., Li, X., Meng, Y., Wu, Z., Guo, M., Zhao, Y., Fu, X. and Han, W. (2012) A facile and specific assay for quantifying microRNA by an optimized RT-qPCR approach. PloS one, 7, e46890.

27. Kernick, K., Woudstra, R., Berjanskii, M., MacKay, S. and Wishart, D.S. (2025) Heatmapper2: web-enabled heat mapping made easy. Nucleic Acids Res, 53, W316–W323.

28. Livak, K.J. and Schmittgen, T.D. (2001) Analysis of relative gene expression data using real-time quantitative PCR and the 2(-Delta Delta C(T)) Method. Methods, 25, 402–408.

29. Kotagama, K., Schorr, A.L., Steber, H.S. and Mangone, M. (2019) ALG-1 Influences Accurate mRNA Splicing Patterns in the Caenorhabditis elegans Intestine and Body Muscle Tissues by Modulating Splicing Factor Activities. Genetics, 212, 931–951.

30. Kato, M., Kashem, M.A. and Cheng, C. (2016) An intestinal microRNA modulates the homeostatic adaptation to chronic oxidative stress in C. elegans. Aging (Albany NY), 8, 1979–2005.

31. Stubna, M.W., Shukla, A. and Bartel, D.P. (2024) Widespread destabilization of Caenorhabditis elegans microRNAs by the E3 ubiquitin ligase EBAX-1. RNA, 31, 51–66.

32. Kent, W.J. (2002) BLAT--the BLAST-like alignment tool. Genome Res, 12, 656–664.

33. Lorenz, R., Bernhart, S.H., Honer Zu Siederdissen, C., Tafer, H., Flamm, C., Stadler, P.F. and Hofacker, I.L. (2011) ViennaRNA Package 2.0. Algorithms Mol Biol, 6, 26.

34. Rishik, S., Hirsch, P., Grandke, F., Fehlmann, T. and Keller, A. (2025) miRNATissueAtlas 2025: an update to the uniformly processed and annotated human and mouse non-coding RNA tissue atlas. Nucleic Acids Res, 53, D129–D137.

35. Brosnan, C.A., Palmer, A.J. and Zuryn, S. (2021) Cell-type-specific profiling of loaded miRNAs from Caenorhabditis elegans reveals spatial and temporal flexibility in Argonaute loading. Nat Commun, 12, 2194.

36. Shaw, W.R., Armisen, J., Lehrbach, N.J. and Miska, E.A. (2010) The conserved miR-51 microRNA family is redundantly required for embryonic development and pharynx attachment in Caenorhabditis elegans. Genetics, 185, 897–905.

37. Alvarez-Saavedra, E. and Horvitz, H.R. (2010) Many families of C. elegans microRNAs are not essential for development or viability. Curr Biol, 20, 367–373.

38. Garcia-Segura, L., Abreu-Goodger, C., Hernandez-Mendoza, A., Dimitrova Dinkova, T.D., Padilla-Noriega, L., Perez-Andrade, M.E. and Miranda-Rios, J. (2015) High-Throughput Profiling of Caenorhabditis elegans Starvation-Responsive microRNAs. PloS one, 10, e0142262.

39. Garcia, S.M., Tabach, Y., Lourenço, G.F., Armakola, M. and Ruvkun, G. (2014) Identification of genes in toxicity pathways of trinucleotide-repeat RNA in C. elegans. Nature structural & molecular biology, 21, 712–720.

40. Kagias, K., Podolska, A. and Pocock, R. (2014) Reliable reference miRNAs for quantitative gene expression analysis of stress responses in Caenorhabditis elegans. BMC Genomics, 15, 222.

41. Blazie, S.M., Geissel, H.C., Wilky, H., Joshi, R., Newbern, J. and Mangone, M. (2017) Alternative Polyadenylation Directs Tissue-Specific miRNA Targeting in Caenorhabditis elegans Somatic Tissues. Genetics, 206, 757–774.

42. Haenni, S., Ji, Z., Hoque, M., Rust, N., Sharpe, H., Eberhard, R., Browne, C., Hengartner, M.O., Mellor, J., Tian, B. et al. (2012) Analysis of C. elegans intestinal gene expression and polyadenylation by fluorescence-activated nuclei sorting and 3’-end-seq. Nucleic Acids Res, 40, 6304–6318.

43. Nakanishi, K. (2024) When Argonaute takes out the ribonuclease sword. J Biol Chem, 300, 105499.

44. Krishnan, K., Steptoe, A.L., Martin, H.C., Wani, S., Nones, K., Waddell, N., Mariasegaram, M., Simpson, P.T., Lakhani, S.R., Gabrielli, B. et al. (2013) MicroRNA-182-5p targets a network of genes involved in DNA repair. RNA, 19, 230–242.

45. Krist, B., Florczyk, U., Pietraszek-Gremplewicz, K., Jozkowicz, A. and Dulak, J. (2015) The Role of miR-378a in Metabolism, Angiogenesis, and Muscle Biology. Int J Endocrinol, 2015, 281756.

46. Rane, S., He, M., Sayed, D., Vashistha, H., Malhotra, A., Sadoshima, J., Vatner, D.E., Vatner, S.F. and Abdellatif, M. (2009) Downregulation of miR-199a derepresses hypoxia-inducible factor-1alpha and Sirtuin 1 and recapitulates hypoxia preconditioning in cardiac myocytes. Circ Res, 104, 879–886.

47. Makeyev, E.V., Zhang, J., Carrasco, M.A. and Maniatis, T. (2007) The MicroRNA miR-124 promotes neuronal differentiation by triggering brain-specific alternative pre-mRNA splicing. Mol Cell, 27, 435–448.

48. Rhim, J., Baek, W., Seo, Y. and Kim, J.H. (2022) From Molecular Mechanisms to Therapeutics: Understanding MicroRNA-21 in Cancer. Cells, 11.

49. Brueckner, B., Stresemann, C., Kuner, R., Mund, C., Musch, T., Meister, M., Sultmann, H. and Lyko, F. (2007) The human let-7a-3 locus contains an epigenetically regulated microRNA gene with oncogenic function. Cancer Res, 67, 1419–1423.

50. Kato, M., Paranjape, T., Muller, R.U., Nallur, S., Gillespie, E., Keane, K., Esquela-Kerscher, A., Weidhaas, J.B. and Slack, F.J. (2009) The mir-34 microRNA is required for the DNA damage response in vivo in C. elegans and in vitro in human breast cancer cells. Oncogene, 28, 2419–2424.

51. Isik, M., Blackwell, T.K. and Berezikov, E. (2016) MicroRNA mir-34 provides robustness to environmental stress response via the DAF-16 network in C. elegans. Sci Rep, 6, 36766.

52. Bommer, G.T., Gerin, I., Feng, Y., Kaczorowski, A.J., Kuick, R., Love, R.E., Zhai, Y., Giordano, T.J., Qin, Z.S., Moore, B.B. et al. (2007) p53-mediated activation of miRNA34 candidate tumor-suppressor genes. Curr Biol, 17, 1298–1307.

53. Spurgers, K.B., Gold, D.L., Coombes, K.R., Bohnenstiehl, N.L., Mullins, B., Meyn, R.E., Logothetis, C.J. and McDonnell, T.J. (2006) Identification of cell cycle regulatory genes as principal targets of p53-mediated transcriptional repression. J Biol Chem, 281, 25134–25142.

54. He, L., He, X., Lowe, S.W. and Hannon, G.J. (2007) microRNAs join the p53 network--another piece in the tumour-suppression puzzle. Nat Rev Cancer, 7, 819–822.

55. Kim, H., Kim, J., Yu, S., Lee, Y.Y., Park, J., Choi, R.J., Yoon, S.J., Kang, S.G. and Kim, V.N. (2020) A Mechanism for microRNA Arm Switching Regulated by Uridylation. Mol Cell, 78, 1224–1236 e1225.

56. Mullokandov, G., Baccarini, A., Ruzo, A., Jayaprakash, A.D., Tung, N., Israelow, B., Evans, M.J., Sachidanandam, R. and Brown, B.D. (2012) High-throughput assessment of microRNA activity and function using microRNA sensor and decoy libraries. Nat Methods, 9, 840–846.

57. Krol, J., Loedige, I. and Filipowicz, W. (2010) The widespread regulation of microRNA biogenesis, function and decay. Nat Rev Genet, 11, 597–610.

58. Mi, S., Cai, T., Hu, Y., Chen, Y., Hodges, E., Ni, F., Wu, L., Li, S., Zhou, H., Long, C. et al. (2008) Sorting of small RNAs into Arabidopsis argonaute complexes is directed by the 5’ terminal nucleotide. Cell, 133, 116–127.

59. Okamura, K., Liu, N. and Lai, E.C. (2009) Distinct mechanisms for microRNA strand selection by Drosophila Argonautes. Mol Cell, 36, 431–444.

60. Kotagama, K., Grimme, A.L., Braviner, L., Yang, B., Sakhawala, R.M., Yu, G., Benner, L.K., Joshua-Tor, L. and McJunkin, K. (2024) Catalytic residues of microRNA Argonautes play a modest role in microRNA star strand destabilization in C. elegans. Nucleic Acids Res, 52, 4985–5001.

61. Stein, L., Mangone, M., Schwarz, E., Durbin, R., Thierry-Mieg, J., Spieth, J. and Sternberg, P. (2001) WormBase: network access to the genome and biology of Caenorhabditis elegans (vol 29, pg 82, 2001). Nucleic Acids Research, 29, 1012.

