## Supplemental Figures for "An AI-Guided Framework Reveals Conserved Features Governing microRNA Strand Selection"

Additional File 1 for:

**This PDF includes:**

**Supplemental Figs. S1-S10**

**Table of Contents**

### Additional File 1

**Fig. S1: Strand Usage per Category in Representative miRNA..…..…3**

**Fig. S2: Supplemental Figure S2: Comparative analysis of miRNA detection and expression profiles across three datasets (1) ..4**

**Fig. S3: Supplemental Figure S2: Comparative analysis of miRNA detection and expression profiles across three datasets (2) ..6**

**Fig. S4: Supplemental Figure S2: Comparative analysis of miRNA detection and expression profiles across three datasets (3) ..8**

**Fig. S5: HiTmiSS Comparison and Intestine Tissue *Singletons/Switchers*……………………………………..……..…10**

**Fig. S6: Testing the Target Stabilization Hypothesis......................…11**

**Fig. S7: Overview of the miRNA Strand Usage Prediction Model…..12**

**Fig. S8: Overview of Model Performance………………………………13**

**Fig. S9: Predicted Strand Usage in *5p* and *3p Singletons*……..……14**

**Fig. S10: Comparative analysis of miRNA strand selection in metazoans…………………………………………………….…..15**

**A**

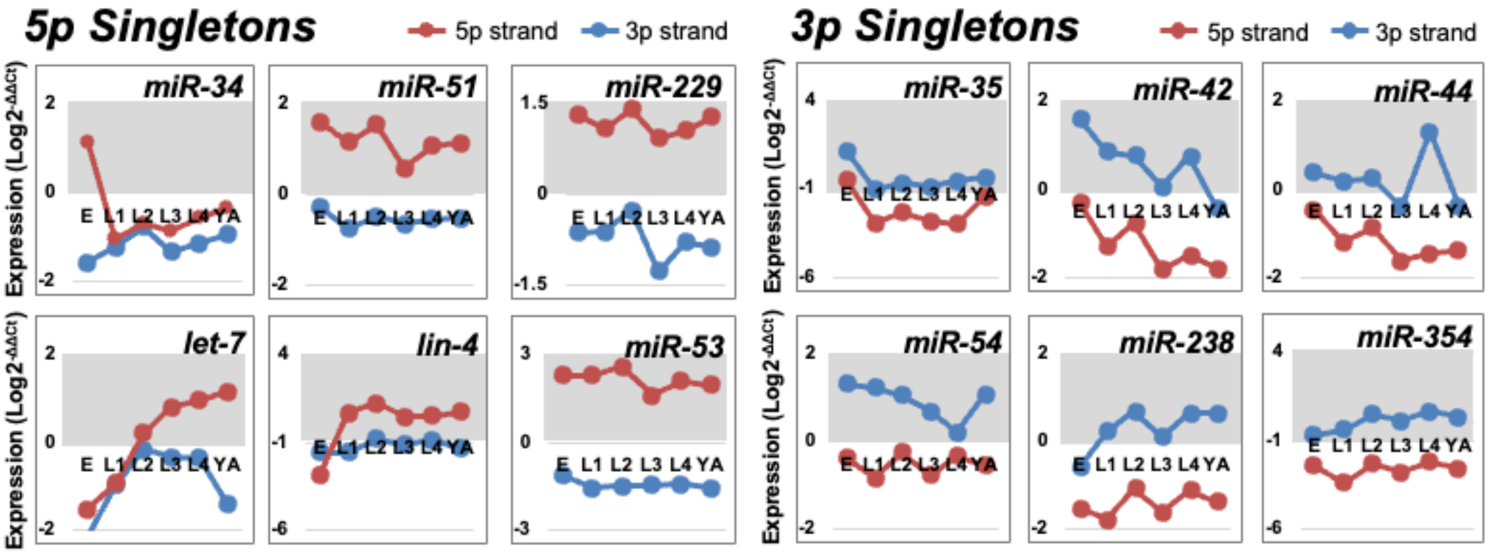


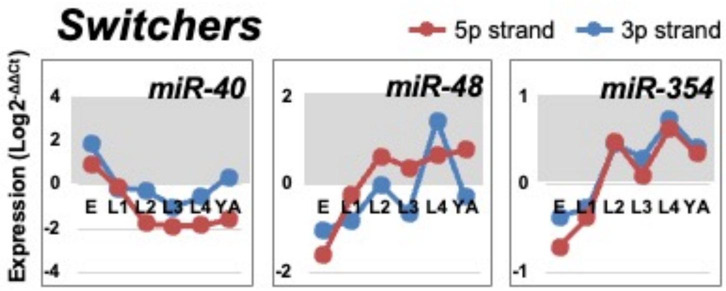


**B**

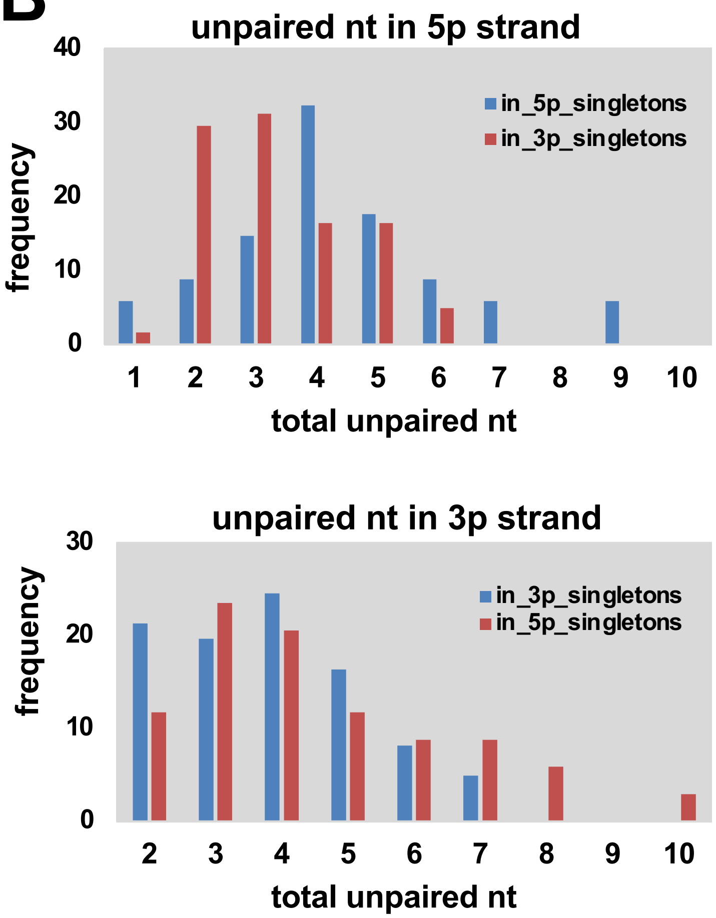


**Supplemental Figure S1: Strand Usage per Category in Representative miRNAs**. (**A**) **Normalized mean expression values (n=3) for the 5p strand (red) and 3p strand (blue) of several representative miRNAs that fall into one of our categories (*5p Singleton*, *3p Singleton*, or *Switcher*).** Expression is demonstrated to fluctuate across developmental time for each of these miRNAs in mixed tissue, following their expression from embryo stage (E) through the four larval stages (L1-L4) and in young adult stage (YA). Within each *Singleton* category we find well characterized canonical regulators of development such as *let-7*, *lin-4*, *miR-35* and *miR-53*. Within the *Switcher* category the trends are more diverse, with some miRNAs like *miR-40* altering which strand is used after embryo stage, while others like *miR-48* display strand switching later in development. Still others like *miR-354* show relatively equal strand usage throughout all developmental stages. (**B**) **Distribution of unpaired nucleotides in miRNA hairpin regions across** (mean unpaired count: 5p strand in *5p Singletons*: 4.32; 5p strand in *3p Singletons*: 3.31; 3p strand in *3p Singletons*: 3.70; 3p strand in *5p Singletons*: 4.32)

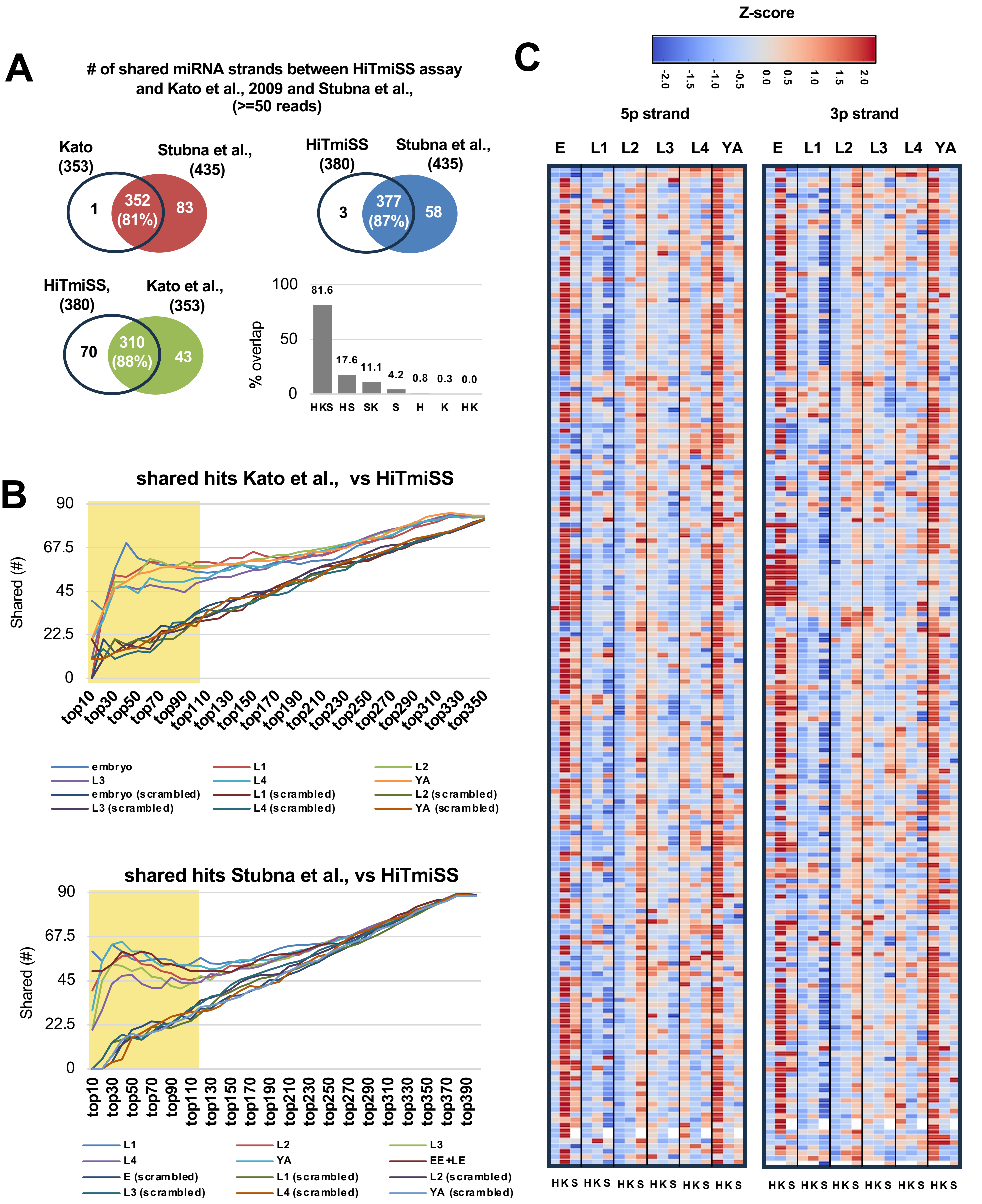
A
### of shared miRNA strands between HiTmiSS assay and Kato et al., 2009 and Stubna et al., (>=50 reads)
Kato (353)
Stubna et al., (435)
1
352 (81%)
83
HiTmiSS (380)
Stubna et al., (435)
3
377 (87%)
58
HiTmiSS, (380)
Kato et al., (353)
70
310 (88%)
43
% overlap
100
50
0
81.6
17.6
11.1
4.2
0.8
0.3
0.0
HKS HS SK S H K HK
B
shared hits Kato et al., vs HiTmiSS
Shared (#)
90
67.5
45
22.5
0
top10 top30 top50 top70 top90 top110 top130 top150 top170 top190 top210 top230 top250 top270 top290 top310 top330 top350
embryo
L1
L2
L3
L4
YA
embryo (scrambled)
L1 (scrambled)
L2 (scrambled)
L3 (scrambled)
L4 (scrambled)
YA (scrambled)
shared hits Stubna et al., vs HiTmiSS
Shared (#)
90
67.5
45
22.5
0
top10 top30 top50 top70 top90 top110 top130 top150 top170 top190 top210 top230 top250 top270 top290 top310 top330 top350 top370 top390
L1
L2
L3
L4
YA
EE+LE
E (scrambled)
L1 (scrambled)
L2 (scrambled)
L3 (scrambled)
L4 (scrambled)
YA (scrambled)
C
Z-score
-2.0 -1.5 -1.0 -0.5 0.0 0.5 1.0 1.5 2.0
5p strand
3p strand
E L1 L2 L3 L4 YA
E L1 L2 L3 L4 YA
HKS HKS HKS HKS HKS HKS
HKS HKS HKS HKS HKS HKS

**Supplemental Figure S2: Comparative analysis of miRNA detection and expression profiles across three datasets: Kato et al. (RNA-seq), Stubna et al. (RNA-seq), and HiTmiSS.** HiTmiSS outperforms RNA-Seq in detecting biologically relevant miRNA strand dynamics. To benchmark the performance of our HiTmiSS assay, we compared our results in **Main Figure 1** with small RNA-seq data from Kato et al., and from Stubna et al., which profiled total small RNA (including miRNAs) across *C. elegans* development.

**(A) Venn diagrams illustrating the overlap in miRNA strand detection between Kato et al., Stubna et al., and HiTmiSS datasets.** *Top left*: Kato et al., and Stubna et al., share 81% of detected miRNA strands. *Top right*: HiTmiSS and Stubna et al., share 87% of strands. *Bottom left*: HiTmiSS and Kato et al., share 88% of strands. *Bottom right*: intersection plot showing all three datasets (H = HiTmiSS, K = Kato et al., S = Stubna et al.,) share 81.6% of unique miRNA strands. These overlaps highlight a strong concordance in miRNA detection across independent methods and platforms.

**(B) Comparative ranking analysis of the most abundant miRNAs between HiTmiSS and Kato et al., (Top) or Stubna et al., (Bottom).** Charts show the overlap of the top 10 to 350 most highly expressed benchmarked against scrambled controls. The substantial overlap demonstrates strong agreement between HiTmiSS and sequencing-based datasets.

**(C) Heatmaps of 5p and 3p miRNA expression profiles across developmental stages in HiTmiSS, Kato, and Stubna.** Stubna and Kato (RNA-seq) show broader contrasts, reflecting a larger dynamic range typical of sequencing, whereas HiTmiSS values are more compressed but exhibit smoother transitions between stages. In all three datasets, stage-to-stage profiles remain consistent: many embryonic miRNAs decrease from blue (low) to red (high) in larval stages, while others peak in young adults (YA), underscoring the preservation of developmental trends across platforms.

**Reference**:

Kato, M., de Lencastre, A., Pincus, Z. & Slack, F. J.
Dynamic expression of small non-coding RNAs, including novel microRNAs and piRNAs/21U-RNAs, during Caenorhabditis elegans development. *Genome Biol* **10**, R54 (2009).

Stubna MW, Shukla A, Bartel DP.
Widespread destabilization of Caenorhabditis elegans microRNAs by the E3 ubiquitin ligase EBAX-1 RNA . 2024 Dec 16;31(1):51-66

#### Supplemental Figure S3

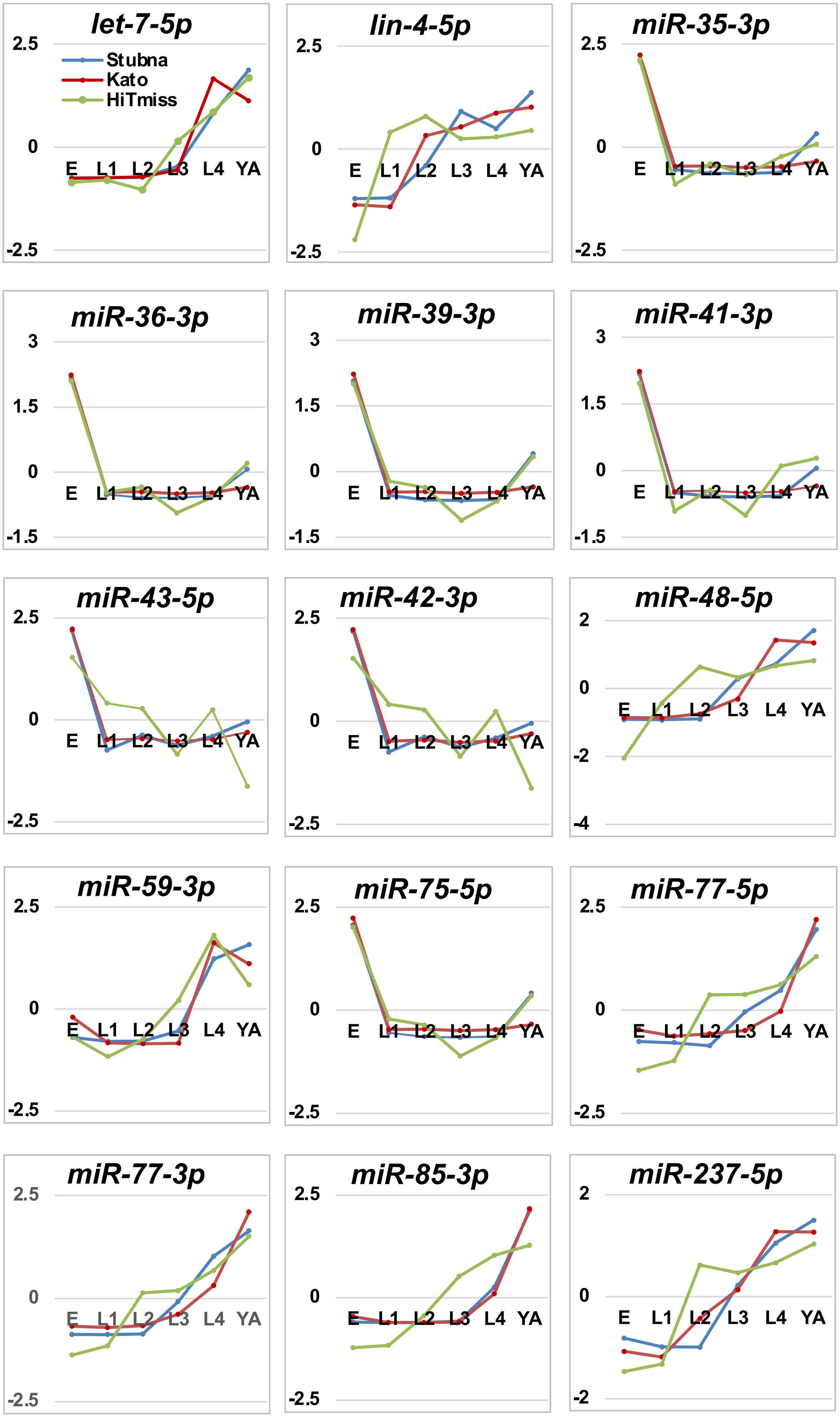

**Supplemental Figure S3: Comparative analysis of miRNA detection and expression profiles across three datasets: Kato et al. (RNA-seq), Stubna et al. (RNA-seq), and HiTmiSS (II).** Representative examples of 15 miRNAs strand usage detected in common across all three datasets, illustrating similar developmental trajectories despite methodological differences.

**Reference**:

Kato, M., de Lencastre, A., Pincus, Z. & Slack, F. J.
Dynamic expression of small non-coding RNAs, including novel microRNAs and piRNAs/21U-RNAs, during Caenorhabditis elegans development. *Genome Biol* **10**, R54 (2009).

Stubna MW, Shukla A, Bartel DP.
Widespread destabilization of Caenorhabditis elegans microRNAs by the E3 ubiquitin ligase EBAX-1 RNA . 2024 Dec 16;31(1):51-66

Supplemental Figure S4

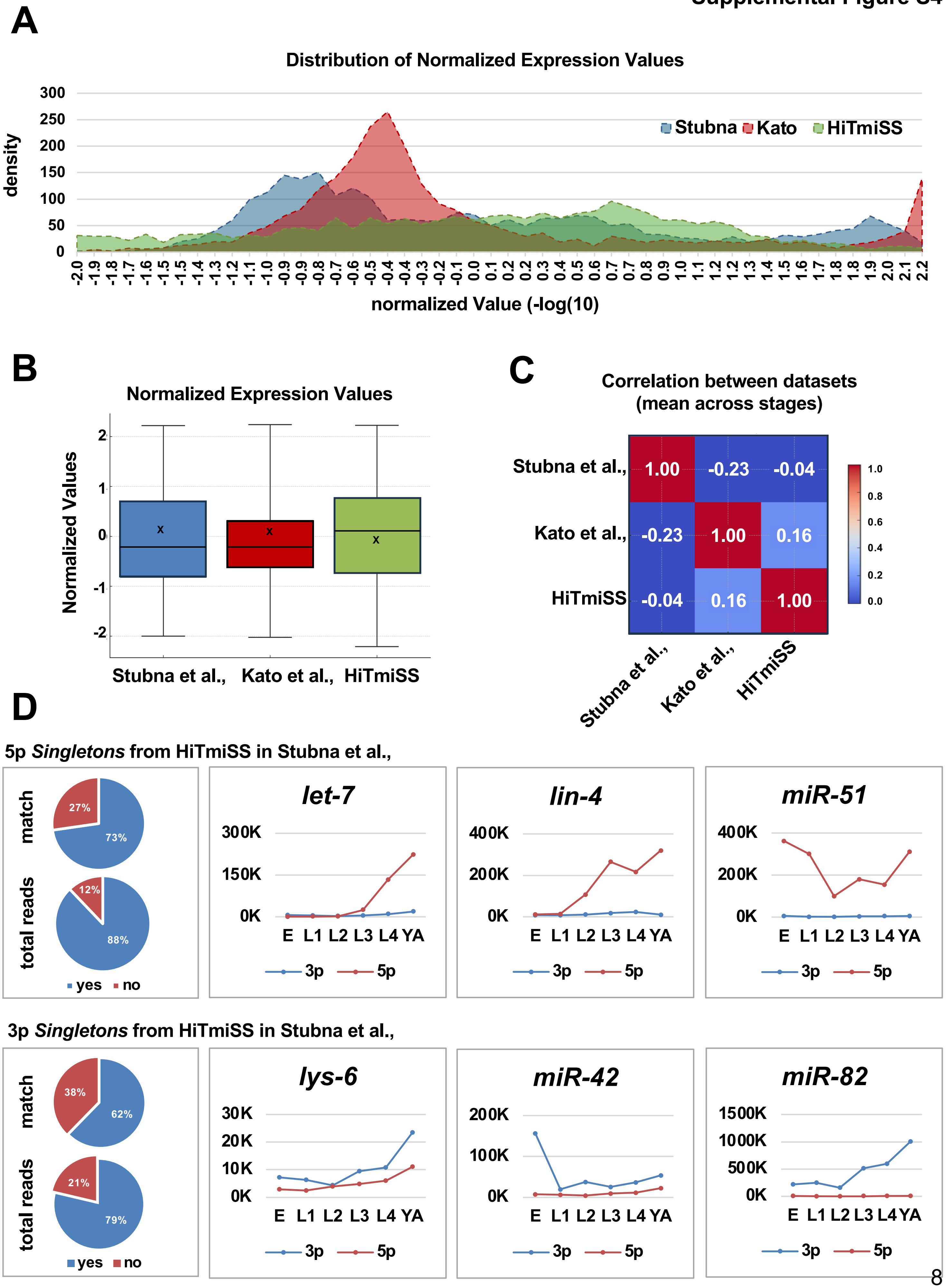

**Supplemental Figure S4. Comparative analysis of miRNA detection and expression profiles across three datasets: Kato et al. (RNA-seq), Stubna et al. (RNA-seq), and HiTmiSS (III).**

**(A) Distribution of normalized expression values.** Density plots illustrate the global distribution of normalized miRNA expression values (−log10) across three independent datasets: Stubna et al. (blue), Kato et al. (red), and HiTmiSS (green). Each dataset shows a distinct profile, with Kato et al. displaying a narrower, higher peak of moderately expressed miRNAs, Stubna et al. showing a broader distribution enriched in lower expression values, and HiTmiSS capturing a wider dynamic range with tails extending to both low and high values. These differences reflect both methodological variation (RNA-seq versus qPCR-based HiTmiSS) and the ability of HiTmiSS to detect a broader spectrum of expression levels.

**(B) Normalized expression values across datasets.** Boxplots summarize the distribution of normalized expression values for each dataset. The central line indicates the median, boxes represent the interquartile range, whiskers extend to 1.5× interquartile range (IQR), and the "x" marks the mean. Although median values are broadly similar, HiTmiSS shows a wider spread, consistent with its greater sensitivity to lowly and highly expressed strands. Stubna et al. and Kato et al. display tighter ranges, highlighting the constraints of RNA-seq–based quantification compared with targeted amplification methods.

**(C) Correlation between datasets.** Heatmap of Pearson correlation coefficients (mean across developmental stages) quantifies the similarity between expression profiles. Stubna et al. and Kato et al. are negatively correlated ($r = -0.23$), suggesting systematic differences in quantification or coverage. HiTmiSS shows weak but positive correlation with Kato et al. ($r = 0.16$), while no meaningful correlation is observed with Stubna et al. ($r = -0.04$). This analysis underscores the unique contribution of HiTmiSS, which provides complementary information to RNA-seq datasets rather than simply replicating them.

**(D) Comparison of individual *Singletons*.** Left: Pie charts summarize the overlap of 5p (top) and 3p (bottom) *Singletons* identified by HiTmiSS with those detected in Stubna et al. The top pies indicate the fraction of cases where HiTmiSS-defined 5p or 3p Singletons were also consistently classified as *Singletons* in Stubna et al. (i.e., one strand shows higher expression than its partner across all developmental stages for a given miRNA). The bottom pies present the same comparison but based on the strand with the highest cumulative read counts across all stages. 73-88% of 5p and 62-79% of 3p identified as *Singletons* by HiTmiSS are also *Singletons* in Stubna et al., Right: Developmental expression profiles of representative 5p (top) and 3p (Bottom) Singletons across embryonic (E) and larval stages (L1–L4) to young adult (YA). Well-characterized developmental regulators 5p Singletons (*let-7* and *lin-4*) show strong induction during larval stages, with 5p strands dominating. *miR-51* and *miR-42* exhibit stable expression phenocopy 5p and 3p Singleton presence. *lys-6* displays gradual upregulation, while *miR-82* reaches very high abundance at later stages. These examples highlight the improved resolution of HiTmiSS in capturing strand-specific expression dynamics.

**Reference**:

Kato, M., de Lencastre, A., Pincus, Z. & Slack, F. J.
Dynamic expression of small non-coding RNAs, including novel microRNAs and piRNAs/21U-RNAs, during Caenorhabditis elegans development. *Genome Biol* **10**, R54 (2009).

Stubna MW, Shukla A, Bartel DP.
Widespread destabilization of Caenorhabditis elegans microRNAs by the E3 ubiquitin ligase EBAX-1 RNA . 2024 Dec 16;31(1):51-66

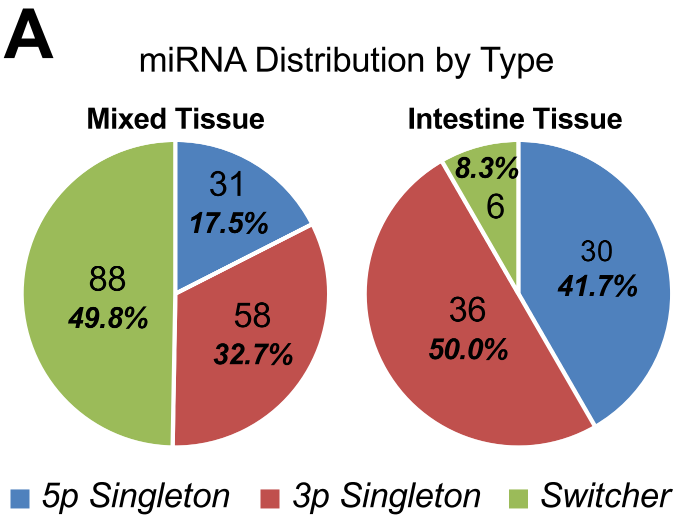


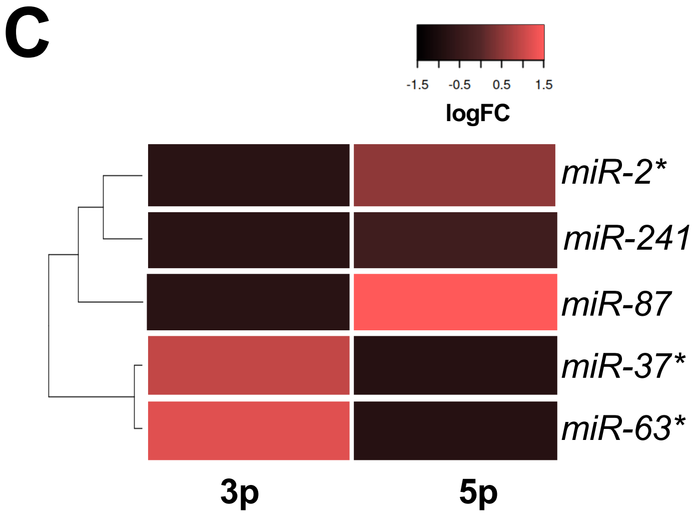


**B**

| miRNA | stage | preferred strands | | HiTmiSS FC Ratio |
|---|---|---|---|---|
| | | ***HiTmiSS*** | ***Brosnan et al.,*** | |
| *miR-2* | Embryo | 5p | 5p | +4.45 |
| *miR-36* | | 3p | 3p | +9.67 |
| *miR-37* | | 3p | 3p | +40.83 |
| *miR-87* | | 5p | 5p/3p | +12.85 |
| *miR-241* | Adult | 5p | 5p/3p | -6.12 |
| *miR-63* | | 3p | 3p | +134.32 |

**Supplemental Figure S5: HiTmiSS Comparison and Intestine Tissue *Switchers*. (A)** Comparison of the distribution of *5p Singletons*, *3p Singletons*, and *Switchers* between HiTmiSS analysis results in mixed tissue (left) and intestine tissue (right). Results expressed as a percentage of total above-threshold miRNAs per analysis. **(B)** List of six identified intestine *Singletons* (embryo or adult) in both our study and Brosnan et al., perferred strand at that stage per HiTmiSS; preferred strand in mixed-stage ALG1 immunoprecipitation per Brosnan et al. (or 5p/3p if neither strand significantly different from N2); Fold change ratio per HiTmiSS. **(C)** Heatmap showing the identification of the six HiTmiSS-identified intestine *Switchers* in the Brosnan et al. dataset. 5p and 3p strand quantity expressed as fold change ALG1 immunoprecipitation vs. N2 (mixed stage).

**Reference:**

Brosnan et al., 2021 (Cell-type-specific profiling of loaded miRNAs from Caenorhabditis elegans reveals spatial and temporal flexibility in Argonaute loading) Nat Commun . 2021 Apr 13;12(1):2194.

**Supplemental Figure S6**

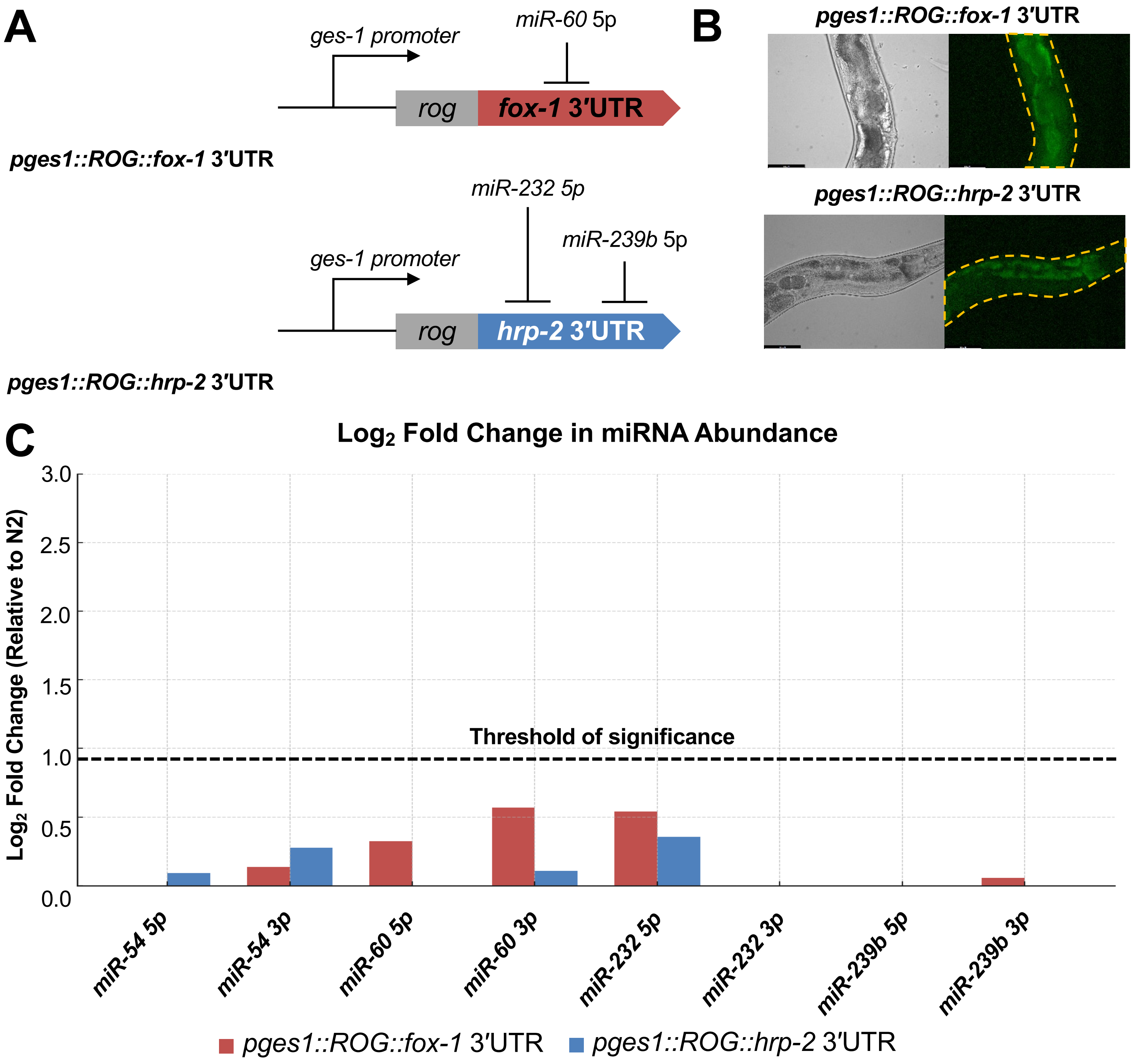


**Supplemental Figure S6: Testing the Target Stabilization Hypothesis**. **(A) Constructs used to overexpress GFP under the control of the *fox-1* 3′UTR (top) or *hrp-2* 3′UTR (bottom) using the promoter for the intestine-specific gene *ges-1.*** miRNA strands which bind each target 3′UTR (shown above) are typically not expressed in wildtype N2 *C. elegans*. **(B) Representative brightfield (left) and GFP (right) images of *C. elegans* strains expressing these constructs, demonstrating fluorescence in the intestine. (C) Change in abundance of each miRNA strand, expressed as $\log_2$ of the difference in mean expression value as per our HiTmiSS assay**. None of the tested miRNA strands were upregulated 2-fold or more ($\log_2 > 1$, displayed as “Threshold of significance) when overexpressing either test 3′UTR. These data suggest that overexpression of targets for a miRNA strand does not lead to retention of that strand, but rather strand selection occurs independent of the presence or absence of target 3′UTRs.

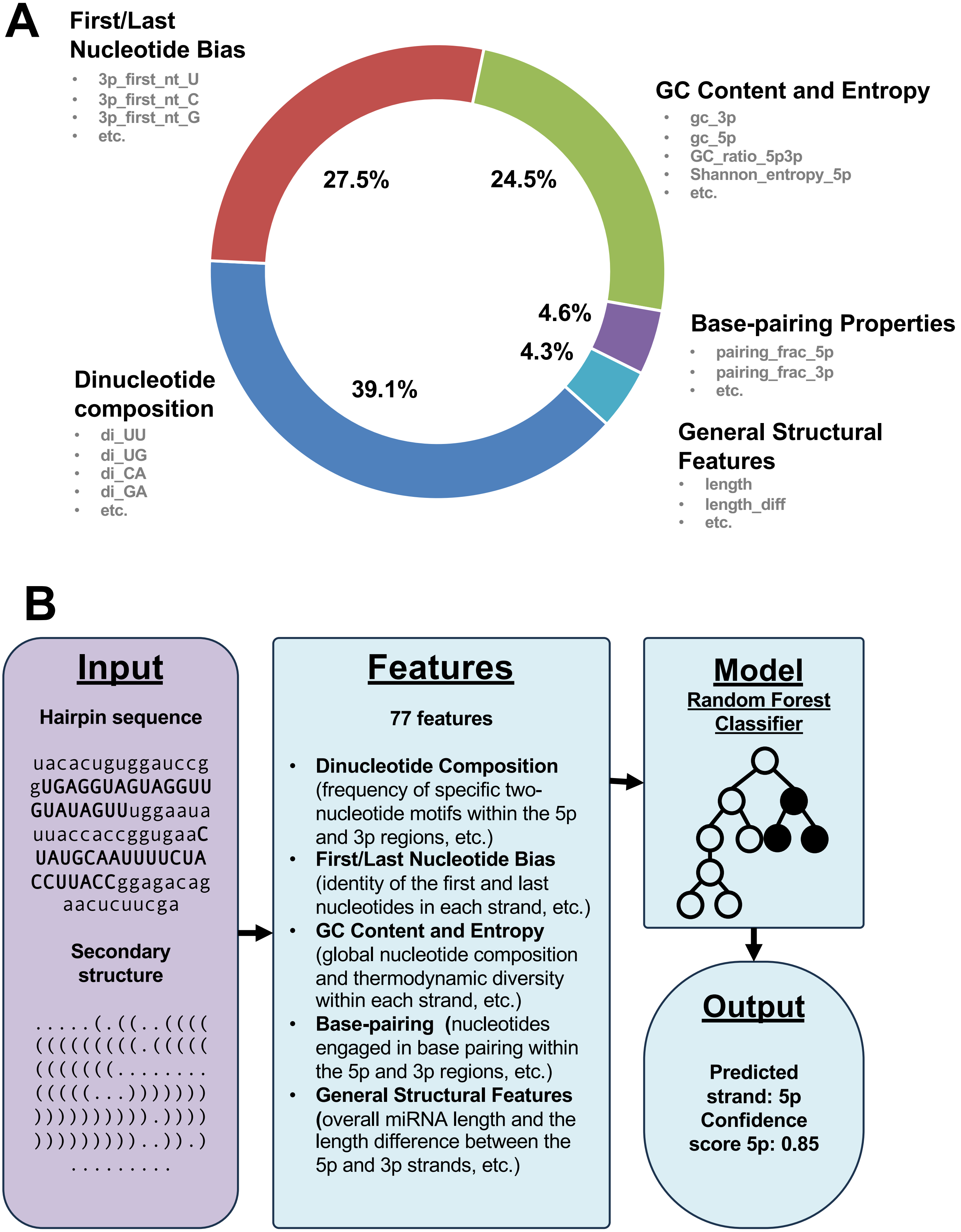


**Supplemental Figure S7: Overview of the miRNA Strand Usage Prediction Model**. **(A)** Weight assigned to each feature of the model. **(B)** Flowchart illustrating the computational pipeline developed to predict 5p or 3p strand usage in miRNA precursors using machine learning. Sample *let-7* input and output shown.

Supplemental Figure S8

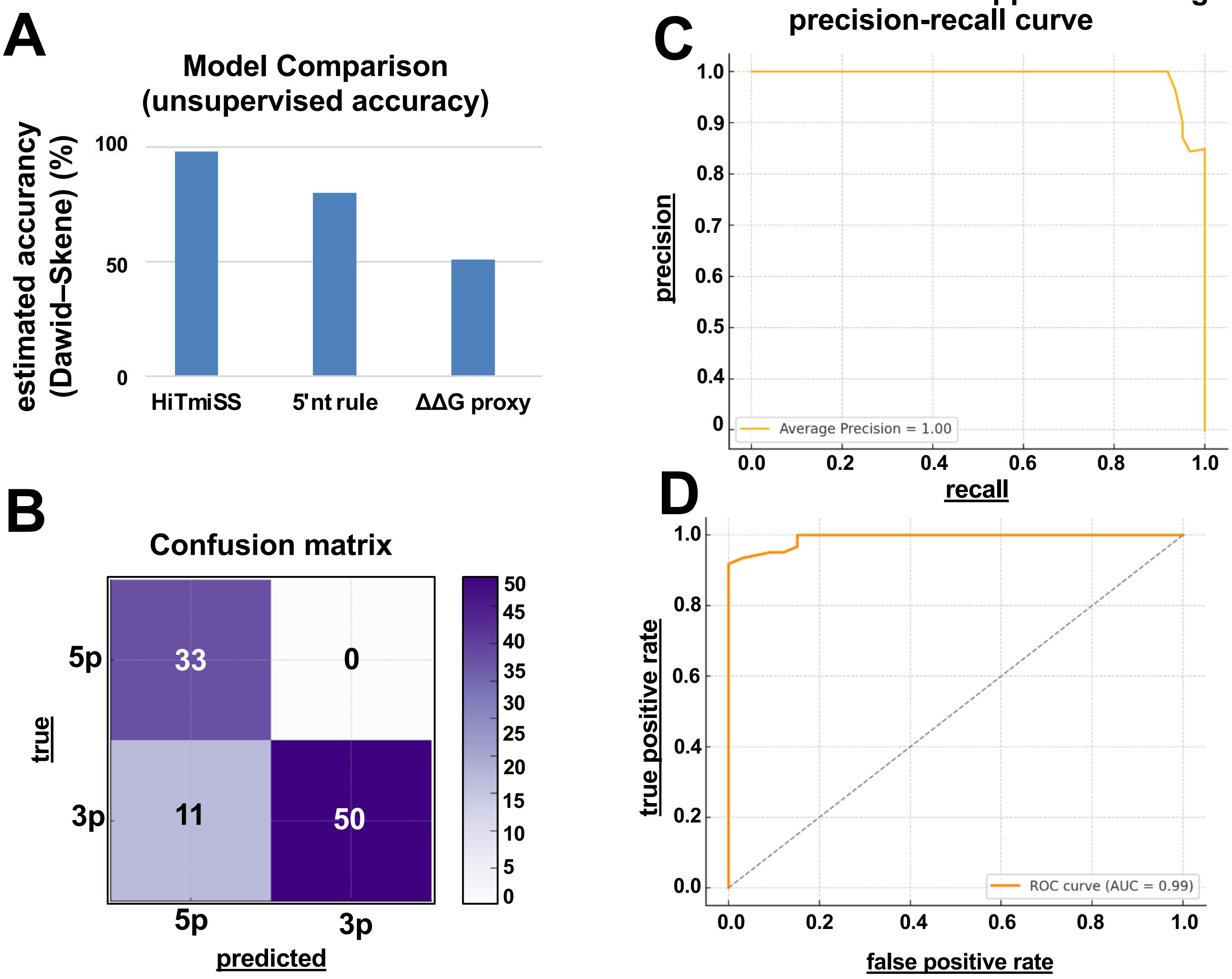


**Supplemental Figure S8: Overview of Model Performance**. **(A)** Bars show the estimated accuracy (prior-weighted diagonal of the inferred 2×2 confusion matrix) for three predictors evaluated on 854 human miRNA duplexes: HiTmiSS, a 5′-nucleotide rule (U>A>C>G), and a ΔΔG end-asymmetry proxy (less-stable 5′ end over the first 5 bp). The Dawid-Skene model jointly infers latent true labels from the three votes and each method's reliability, enabling comparison without a gold standard; higher bars indicate better fit to the joint voting pattern. **(B)** Confusion matrix of the strand usage prediction model. Summarizes the classification performance of the Random Forest model trained on *C. elegans* miRNA data, showing the number of correctly (33 and 50) and incorrectly (0 and 11) predicted 5p and 3p strands. **(C)** Precision-recall curve illustrating the relationship between precision (positive predictive value) and recall (sensitivity) for classifying 5p versus 3p strand usage in miRNA precursors using the Random Forest model. Each point on the curve represents a different decision threshold. The shape of the curve reflects the model's ability to maintain high precision while recovering true positives across a range of thresholds. **(D)** Receiver Operating Characteristic curve of the strand usage prediction model. The curve shows the trade-off between true positive rate (sensitivity) and false positive rate (specificity) for predicting 5p versus 3p strand usage using a Random Forest model trained on *C. elegans* miRNA data. Each point represents a different classification threshold. The area under the curve provides a summary measure of the model's overall performance, with higher values indicating stronger discrimination between 5p and 3p strands. The curve demonstrates reliable strand separation across a range of thresholds.

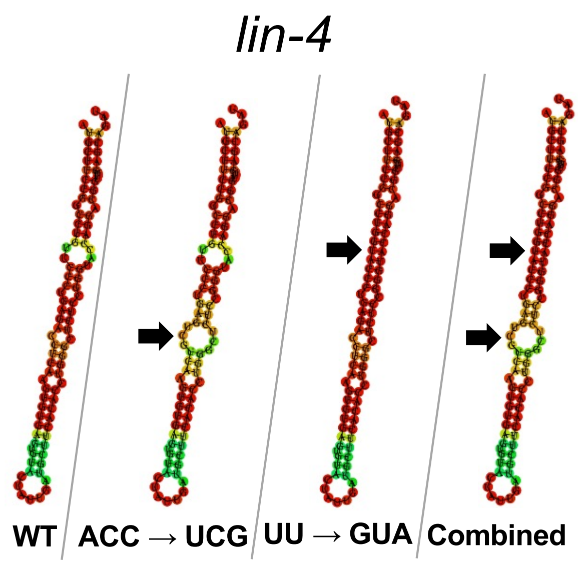


| Variant | 5p | 3p |
|---|---|---|
| WT | 0.81 | 0.19 |
| ACC → UCG | 0.74 | 0.26 |
| UU → GUA | 0.77 | 0.23 |
| Combined | 0.76 | 0.24 |

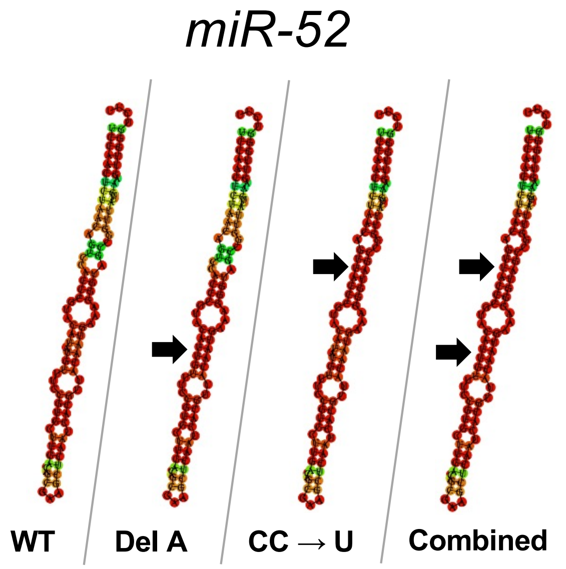


| Variant | 5p | 3p |
|---|---|---|
| WT | 0.89 | 0.11 |
| Del A | 0.83 | 0.17 |
| CC → U | 0.90 | 0.10 |
| Combined | 0.88 | 0.12 |

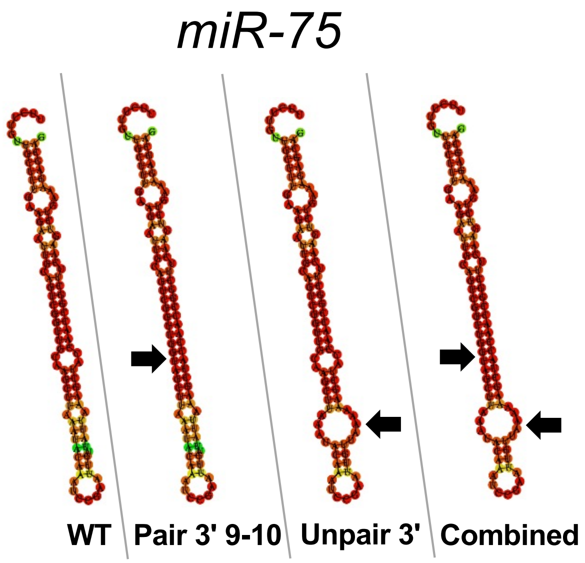


| Variant | 5p | 3p |
|---|---|---|
| WT | 0.36 | 0.64 |
| Pair 3' 9-10 | 0.30 | 0.70 |
| Unpair 3' | 0.60 | 0.40 |
| Combined | 0.57 | 0.43 |

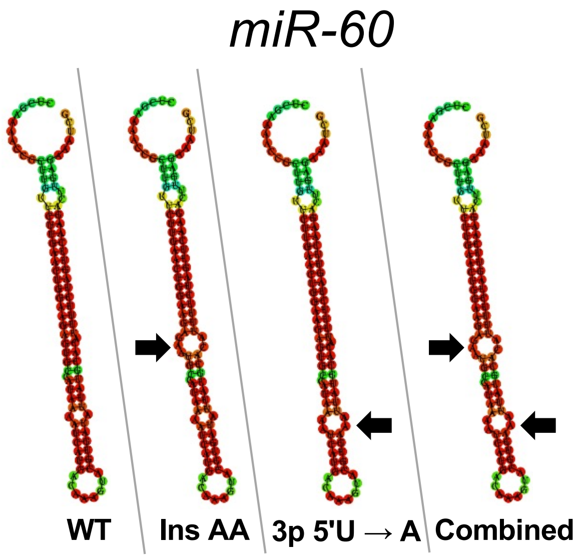


| Variant | 5p | 3p |
|---|---|---|
| WT | 0.41 | 0.59 |
| Ins AA | 0.46 | 0.54 |
| 3p 5'U → A | 0.46 | 0.54 |
| Combined | 0.52 | 0.48 |

**Supplemental Figure S9: Predicted Strand Usage in *5p* and *3p Singletons*.** Representative, well-characterized *5p Singletons* (top) and *3p Singletons* (bottom) were selected and their strand preference predicted by our ML model (WT, far left). Each underwent two *in silico* site-directed mutations, one in the bulge region (middle left) and one in the pairing/identity of the first nucleotide (middle right). Finally, a combination mutation of both is shown (far right). The confidence of prediction is displayed as a decimal, both *5p Singleton* confidence (5p) or *3p Singleton* confidence (3p). These mutations consistently altered predicted strand confidence, with a more pronounced effect in *3p Singletons*.

### Supplemental Figure S10

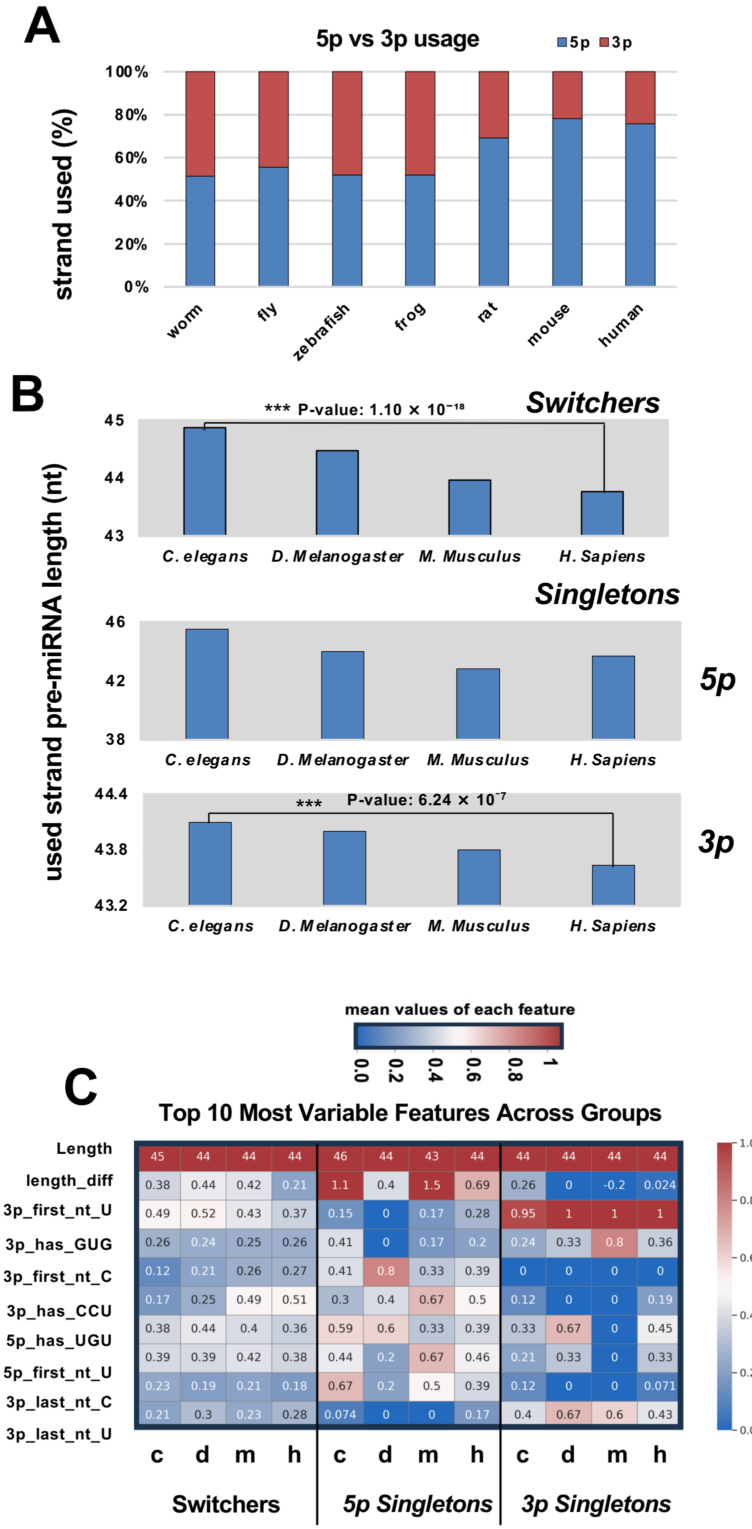


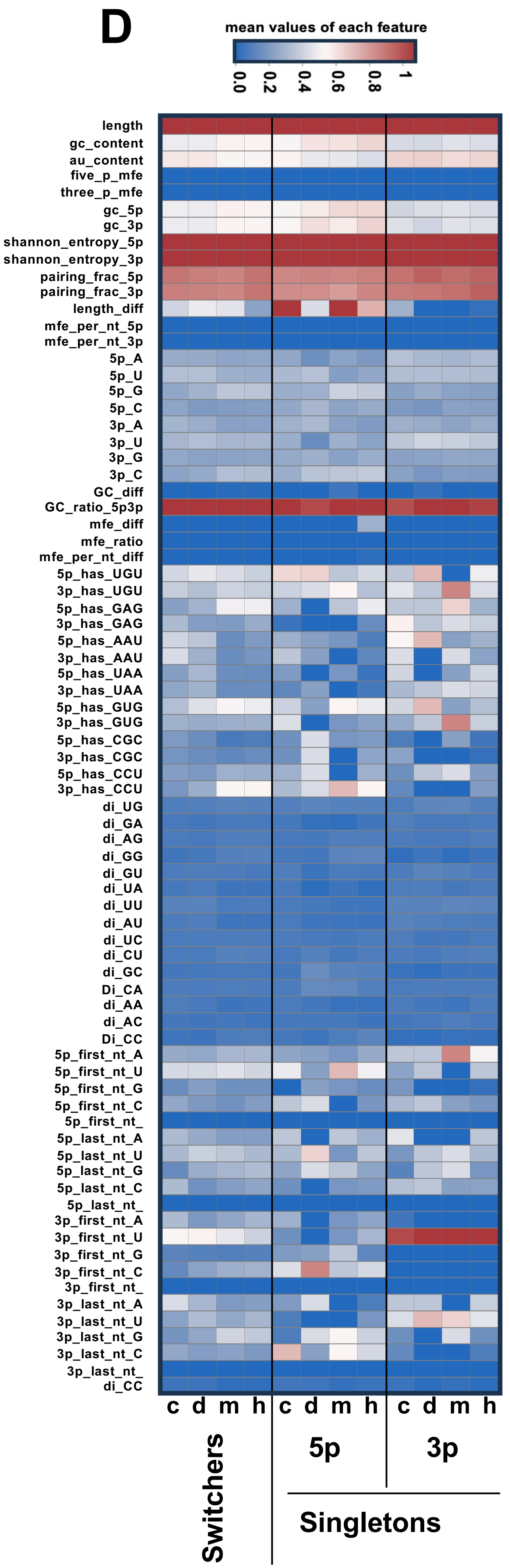


**Supplemental Figure S10: Comparative analysis of miRNA strand selection in metazoans.** (**A**) Top 10 Most Variable Features in *Switchers*, *5p-* and *3p-Singletons*. C = *C. elegans*; *D. melanogaster*; m: *Mus musculus*; h: *homo sapiens.* (**B**) 3p vs 5p strand usage in worm, fly, zebrafish, frog, rat, mouse and human. (**C**) miRNA strand usage length between worm (w), fly (d), mouse (m) and human (h) for miRNA *Switchers*, *5–* and *3p- Singletons.* (**D**) Heatmaps showing the predicted 77 features used in our ML model for worm (w), fly (d), mouse (m) and human (h) for miRNA *Switchers*, *5–* and *3p- Singletons.*
